# Noninvasive profiling of input-output excitability curves in human prefrontal cortex

**DOI:** 10.1101/2025.09.26.678876

**Authors:** Nai-Feng Chen, Umair Hassan, Jessica M. Ross, Lily Forman, James W. Hartford, Juha Gogulski, Sara Parmigiani, Jade Truong, Corey J. Keller, Christopher C. Cline

**Author notes:** Correspondence: Christopher C. Cline, PhD Stanford University Department of Psychiatry and Behavioral Sciences 401 Quarry Road Stanford, CA 94305-5797. Denotes co-senior authorship (Corey J. Keller, Christopher C. Cline).

## Abstract

**Background:** The prefrontal cortex plays a critical role in cognitive control and behavior, and its dysfunction has been linked to numerous psychiatric and neurological disorders. However, noninvasive measurement of prefrontal activity remains challenging, limiting our understanding of how to optimize prefrontal treatments. Input-output relationships reveal how neural circuits respond to different inputs and are essential for determining optimal treatment parameters and understanding individual variability in treatment response, yet systematic investigation of prefrontal input-output relationships has been lacking.

**Objective:** To characterize human prefrontal excitability with input-output (I/O) curves.

**Methods:** We employed transcranial magnetic stimulation (TMS) with electroencephalography in a randomized mixed-block design with 28 healthy participants receiving single-pulse TMS to left dorsolateral prefrontal cortex (dlPFC) across 12 stimulation intensities (60-140% of resting motor threshold). We quantified prefrontal excitability using early local TMS-evoked potentials (EL-TEPs), cortical responses measured locally 20-60 ms post-stimulus.

**Results:** We observed a strong effect of TMS intensity on prefrontal EL-TEP amplitudes. Sigmoidal EL-TEP I/O curves were observed in 57% of participants, with the sigmoidality partially explained by EL-TEP signal quality. Correlations were observed between EL-TEP and motor-evoked potential curve parameters, but intensity parameterization approaches did not significantly differ in explaining inter-individual EL-TEP response variability. Reliable EL-TEPs could be obtained using fewer TMS pulses at high intensities, and test-retest assessments revealed robust I/O curve profiles.

**Conclusions:** These findings provide a systematic noninvasive characterization of prefrontal input-output physiology, establishing a framework for estimating prefrontal excitability. The comparison of various intensity parameterizations motivates the need for enhanced models and individualized measurement of stimulation responses.

**Highlights:**

- We present noninvasive input-output curves for prefrontal TMS.
- EL-TEPs exhibit robust dose-dependent responses to TMS intensity.
- Sigmoidal I/O curves observed in 57% of participants, with a strong dependence on signal quality.
- Correlations observed between MEP and prefrontal EL-TEP curve parameters.
- High test-retest reliability and rapid protocols at ≥110% rMT

## 1. Introduction

The prefrontal cortex plays a critical role in higher-order cognitive functions, emotional regulation, and decision-making processes [1–4]. Its complex physiology underlies fundamental aspects of human behavior and cognition, yet our ability to study its function noninvasively in humans remains limited [5]. Understanding prefrontal cortex physiology is particularly important given that its dysfunction is linked to numerous psychiatric disorders [6–8] and prefrontally targeted treatments such as transcranial magnetic stimulation (TMS) show promise in treating these conditions [9–11]. While these treatments have demonstrated clinical efficacy, response variability points to our incomplete understanding of prefrontal physiology [12–14]. An improved characterization of prefrontal excitability would advance our fundamental knowledge of prefrontal brain function and enhance therapeutic approaches for prefrontal-mediated disorders [15,16].

A fundamental component of neurophysiology studies lies in understanding the relationship between input parameters (e.g., stimulation intensity) and the output of a given brain region. This relationship, often referred to as the input/output (I/O) curve or dose-response curve, provides a comprehensive view of the region’s physiological properties and allows for the systematic examination of its excitability [17]. In the context of TMS treatments, characterizing I/O curves for the prefrontal cortex could help clinicians determine optimal stimulation parameters for individual patients, potentially improving treatment outcomes [18]. In recent years, the utilization of I/O curves has gained recognition as an important method for evaluating human corticospinal excitability through TMS. When specifically examining motor evoked potentials (MEPs), I/O curves typically follow a sigmoidal shape, progressing from no response at low intensities to a plateau at high intensities, with the steepest slope occurring around motor threshold [19]. The characteristics of these curves, including their slope and midpoint, can provide valuable insights into cortical excitability, making them useful biomarkers for assessing disease and treatment effects [20]. While previous studies have extensively characterized I/O curves in the motor cortex [21,22] a comprehensive characterization of I/O curves for estimating prefrontal excitability in humans has been lacking. This gap in knowledge has hindered our understanding of this critical brain region’s physiological properties and the potential for developing targeted neuromodulatory interventions.

TMS combined with electroencephalography (TMS-EEG) has emerged as a powerful tool for noninvasive causal mapping of targeted brain regions with millisecond resolution at the individual participant level. TMS-evoked potentials (TEPs) capture these brain responses and have been used to monitor various neurophysiological states and track changes induced by neuromodulatory treatments [23–25]. Of particular interest for prefrontal cortex studies is the early local TEP (EL-TEP), a short-latency (20-60ms) neural response recorded locally after TMS pulses. The EL-TEP, also referred to as the P20/N40 complex, typically consists of a local positive response at ∼25ms and a negative response at ∼45ms, and is observed when stimulating the dlPFC [26,27] as well as other brain regions [28–30]. EL-TEPs offer several advantages for characterizing prefrontal excitability. They are less affected by off-target sensory confounds compared to later TEP components (>100ms) [31], and recent work has demonstrated they are highly reliable, particularly when stimulating specific prefrontal targets [27]. Additionally, prefrontal EL-TEPs vary significantly as a function of TMS target across the dlPFC [26], and real-time visualization tools have been developed to maximize their detection while minimizing artifacts [32,33]. These properties position the EL-TEP as an ideal measure for characterizing prefrontal I/O relationships and as a potential clinically-relevant biomarker for assessing prefrontal excitability changes induced by neuromodulatory interventions.

In this study, we investigated the input-output curves of prefrontal EL-TEPs. We hypothesized that prefrontal activation, as indexed by EL-TEPs, would exhibit a sigmoidal relationship with increasing TMS intensities, reflecting the nonlinear excitability of cortical physiology. Our findings establish EL-TEPs as a reliable measure of prefrontal excitability and provide a foundation for optimizing individualized brain stimulation treatments.

## 2. Methods

### 2.1 Participants

34 healthy participants (23-63 years old, mean=43.06, SD=12.31, 14 females) initially provided written informed consent under a protocol approved by the Stanford University Institutional Review Board. All participants completed an online screening questionnaire with inclusion criteria as follows: aged 18-65, fluent in English, able to travel to study site, and fully vaccinated against COVID-19. Participants were also required to complete the Quick Inventory of Depressive Symptomatology (16-item, QIDS) self-report questionnaire and were excluded if they scored 11 or higher, indicating moderate or more severe depression. Additional criteria for exclusion included a lifetime history of psychiatric or neurological disorder, substance abuse or dependence in the past month, recent heart attack (<3 months), pregnancy, the presence of contraindications for rTMS, or history of psychotropic medication use. All 34 eligible participants were scheduled for two study visits on separate days to first obtain structural MRI and then complete a TMS-EEG session. All enrolled participants completed an MRI pre-examination screening form provided by Richard M. Lucas Center for Imaging at Stanford University. Six participants were unable to complete the TMS-EEG session and/or were excluded from the analysis; three participants were unable to complete the TMS-EEG session due to equipment issues, two participants were excluded due to data quality issues, and one participant withdrew due to intolerance of the TMS. In total, 28 healthy participants (23-64 years old, mean=43.4, SD=13.1, 13 females) completed a full experiment session (MRI and a TMS-EEG session) and were included in the analysis. Among these 28 participants, 13 (29-64 years old, mean=46.7, SD=12.1, 5 females) participated in a reliability substudy, wherein participants completed a second TMS-EEG session spaced at least one week apart. A table with additional demographic information for participants is available in Table S1.

### 2.2 Transcranial magnetic stimulation

Biphasic single pulses of TMS were delivered using a MagPro X100 stimulator (MagVenture, Denmark). Initial motor hotspotting was performed using a hand-held MagVenture C-B60 figure-of-eight coil (MagVenture, Denmark). The optimal motor hotspot was defined as the coil position from which TMS produced the largest and most consistent twitch in the relaxed right first dorsal interosseous (FDI) muscle, as measured by electromyography (EMG). After hotspotting, a MagVenture Cool-B65 A/P figure-of-eight coil (MagVenture, Denmark) was used, held and positioned automatically by a robotic arm (TMS Cobot, Axilum Robotics, France). To estimate resting motor threshold (rMT), pulses were delivered to the motor hotspot with the coil held tangentially to the scalp and at 45° from midline. rMT was determined to be the minimum intensity that elicited a twitch with amplitude ≥ 50 µV in relaxed FDI in ≥ 5 out of 10 pulses.

Tracking of coil position relative to the participant’s head and MRI data was performed with our custom neuronavigation software, NaviNIBS [34]. Six degree-of-freedom (DOF) poses of the stimulation coil, participant’s head, and stylus tool were tracked via passive infrared optical markers and an array of 3 infrared cameras (OptiTrack PrimeX 41, NaturalPoint, Inc. Corvallis, OR) with motion capture software Motive (v3.0.3, NaturalPoint, Inc. Corvallis, OR) streaming data via IGTLink [35,36] to NaviNIBS. Registration of participant head tracker position was performed based on fiducials at the nasion and near the left and right preauricular points, with scalp-based refinement; re-registration was performed relative to washable marks on the skin corresponding to fiducial locations identified at the start of each session [34]. Anatomical head models were created using SimNIBS headreco [37,38], and MNI targets were transformed into an individual’s native MRI space by a nonlinear inverse deformation field (generated by headreco), with a planned coil orientation generated to position the coil perpendicular to the nearest point on the scalp mesh. This included initial positions for the primary motor cortex (M1) target (MNI -37.3, −18.6, 65.7) as well as the dlPFC target (MNI -37, 23, 47) [26], optimized during hotspotting and artifact minimization steps (see below), respectively. The dlPFC target was selected based on our previous work [26] and its clinical relevance in TMS treatment for depression. Coil position was visualized relative to the gray-matter and scalp surfaces generated by SimNIBS, with real-time metrics of coil alignment distance and angle offsets.

To minimize muscle-related artifacts, we performed our TMS coil angle optimization procedure [33]. To do so, we delivered all single TMS pulses at the predefined initial dlPFC location while maintaining the coil tangential to the scalp and centered on the initial target.

Using NaviNIBS to control the robotic arm [34], we set up automated sampling intervals to systematically vary the coil angle in the coil’s horizontal plane. For initial coil angle optimization, three single pulses of TMS were delivered approximately every 10 degrees between -90 and +90 degrees around the initial target, with the stimulation intensity set at 110% of rMT for all participants. Angle variations were constrained by neuronavigation camera tracking, robot kinematics, and coil cable length limitations. We defined the *early artifact* as the global mean field amplitude (GMFA) averaged between 10-20 ms and identified the coil angle that minimized this artifact. The angle producing the smallest early artifact was then used for a subsequent ramp in intensity up to 140% of rMT (or to the highest tolerable level below that) and a refining angle sweep approximately every 2.5 degrees between −15 and +15 degrees around the new target. The angle producing the smallest early artifact during this process was then selected as the optimal angle and used as the prefrontal target for the duration of the study.

### 2.3 Electroencephalography

EEG data were recorded with a 64-channel TMS-compatible EEG cap with active electrodes (ActiCap, Brain Products GmbH, Germany) and a TMS-compatible amplifier (ActiCHamp plus, Brain products GmbH, Germany) using an acquisition rate of 25 kHz. Electrodes were arranged in a standard montage labeled according to the extended 10- 20 international system. Electrodes were referenced online to Cz and impedances were generally kept below 5 kΩ, with frequent impedance checks. To minimize off-target auditory and somatosensory potentials, masking white noise was delivered through earbuds and additional noise damping was provided by passive earmuffs, as previously described [31].

### 2.4 Electromyography

Motor evoked potentials (MEPs) were recorded from the relaxed first dorsal interosseous (FDI) muscle of the right hand to evaluate corticospinal excitability using electromyography (EMG). Bipolar surface electrodes were placed on the FDI muscle’s belly and the lateral side of the proximal interphalangeal joint of the same finger, and a ground electrode on the corresponding wrist. Prior to electrode placement, the skin was abraded and cleaned to improve EMG signal quality, and the electrodes were secured with medical tape. Single-pulse transcranial magnetic stimulation (TMS) was then applied to the left motor cortex to elicit MEPs in the FDI muscle, and the resulting peak-to-peak MEP amplitudes were recorded. Participants were instructed to maintain a stationary and relaxed posture throughout the experiment.

### 2.5 Experimental design

#### 2.5.1 TMS study design

The study employed a randomized mixed-block design to investigate the I/O relationship of prefrontal excitability. Participants underwent 13 stimulation blocks: 10 targeting dlPFC and 3 targeting M1. To capture a wide range of physiological responses while ensuring participant safety, stimulation intensities were individualized based on each participant’s rMT and tolerability. For dlPFC stimulation, 12 intensities were administered to all participants ranging from 60% to 140% of rMT. The majority of participants (24/28) received stimulations at 60%, 70%, 80%, 90%, 100%, 110%, 115%, 120%, 125%, 130%, 135%, and 140% rMT (Figure 1A) and the other 4 participants received lowered intensities at each step due to tolerability or maximum stimulator output but no conditions were administered <60%. Each dlPFC block included samples of all 12 intensities, grouped into 24 sub-blocks to control for fluctuations in cortical excitability over time. Each sub-block included 10 consecutive single TMS pulses at a single intensity, resulting in 200 dlPFC trials per intensity per session. Intensities were pseudorandomized across sub-blocks, with the constraint that consecutive sub-blocks did not differ by more than 30% in intensity. The inter-stimulus interval (ISI) was jittered between 1.8 and 2.2 seconds (mean = 2 seconds) to mitigate anticipatory effects.

**Figure 1.**
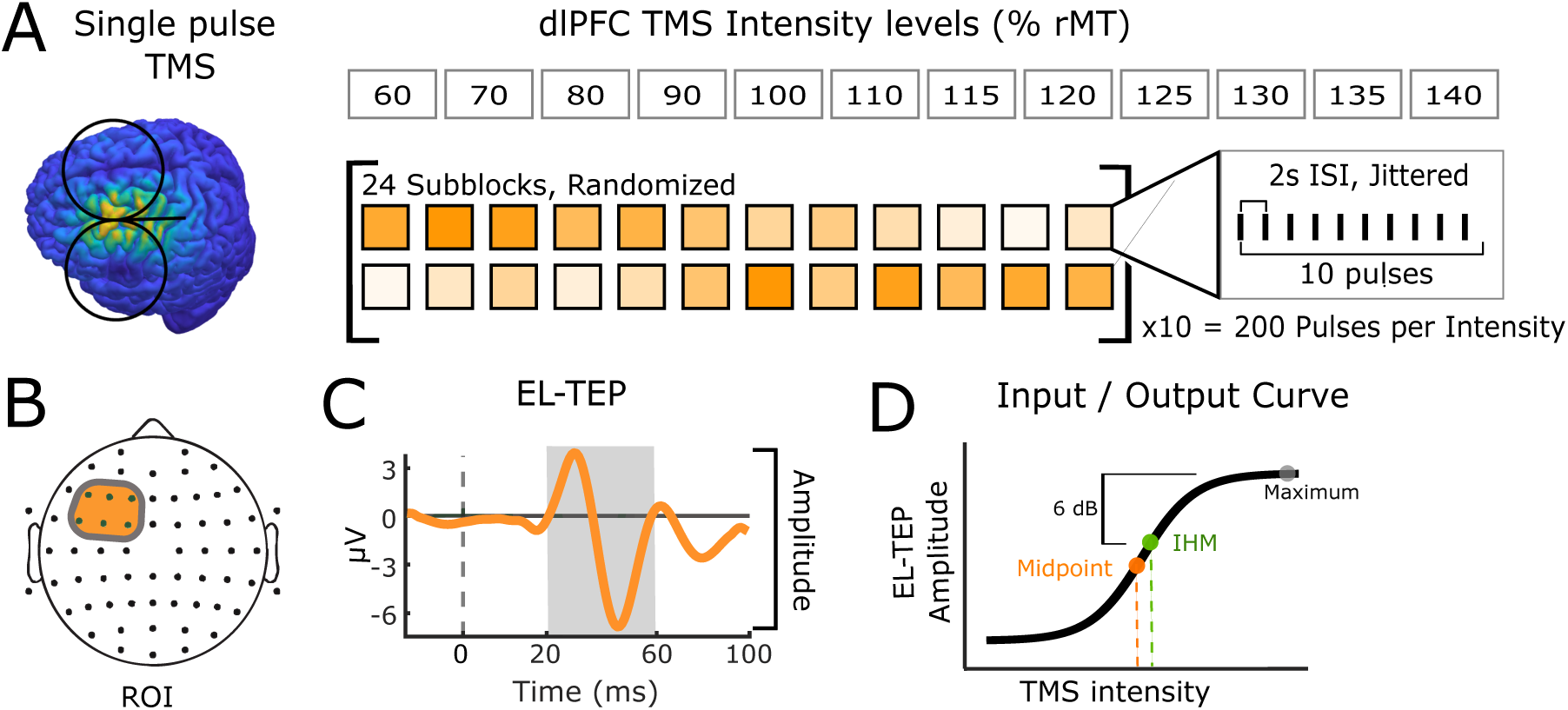
Experimental design and analysis of EL-TEP I/O curves. (A) Single pulse TMS was applied to the dlPFC at twelve intensity levels (60-140% of resting motor threshold, rMT), administered in pseudorandomized order. The inter-stimulus interval (ISI) was jittered around 2 seconds. (B) Region of interest (ROI) on the EEG electrode montage used for EL-TEP quantification. The ROI consists of six electrodes: F5, F3, F1, FC5, FC3, and FC1. (C) Representative EL-TEP waveform showing the time window (20-60 ms, shaded area) used for peak-to-peak amplitude measurement. (D) Schematic representation of the input-output curve. EL-TEP amplitudes were plotted against TMS intensities and fitted with a sigmoidal function, with the midpoint highlighted as a key parameter of the fitted curve.

M1 stimulation followed a similar protocol but with fewer intensity levels. Participants received either 6 (*n* = 11) or 8 (*n* = 17) stimulation intensities, ranging in 10% increments from 80% to 130% or from 70% to 140% rMT respectively, limited by individual tolerability. Each M1 block contained 12 or 16 sub-blocks, maintaining the structure of 10 consecutive pulses per intensity level, yielding 60 trials per intensity. The pseudorandomized intensity distribution and 30% maximum difference between consecutive intensities were maintained. For M1 stimulation, the ISI was jittered between 2.7 and 3.3 seconds (mean = 3 seconds) to reduce intertrial variability of corticospinal responses while minimizing data collection duration [39].

### 2.6. Data Processing

#### 2.6.1. Preprocessing of TMS-EEG data

dlPFC TEPs were preprocessed using version 2.0.1 of the fully automated AARATEP pipeline [40], batched by intensity within each session. Epochs were extracted from 900 ms before to 900 ms after each TMS pulse. Data between 2 ms before to 12 ms after each pulse were replaced with values interpolated by autoregressive extrapolation and blending, downsampled to 1 kHz, and baseline-corrected based on mean values between 500 to 10 ms before the pulse. Epochs were then high-pass filtered above 1 Hz with a modified filtering approach to reduce the spread of pulse-related artifact into baseline time periods. Bad channels were rejected via quantified noise thresholds and replaced with spatially interpolated values (mean ± standard deviation of 1.67±1.68 channels rejected in this data). Per-channel averages were calculated with trimmed mean (outliers of 10% were removed). Eye blink artifacts were attenuated by a dedicated round of independent component analysis (ICA) and eye-specific component labeling and rejection using ICLabel [41] (1.96±0.85 components rejected), a modification from the original AARATEP pipeline introduced in version 2. Various non-neuronal noise sources were attenuated with SOUND [42,43]. To quantify early evoked muscle artifact (12-20 ms), we applied this partial-intermediate preprocessing. Then, artifacts up to 20 ms were reduced via a specialized decay fitting and removal procedure. Line noise was attenuated with a bandstop filter between 58-62 Hz (3rd order). Additional artifacts were attenuated with a second stage of ICA and ICLabel labeling and rejection, with rejection criteria targeted at removing any clearly non-neural signals (65.60%±9.56% of components, representing 43.71%±18.88% variance at this stage). Data were again interpolated between -2 and 12 ms with autoregressive extrapolation and blending, low-pass filtered below 100 Hz, and average re-referenced. For complete details of the pipeline implementation, see Cline et al. [40] and source code at github.com/chriscline/AARATEPPipeline.

#### 2.6.2 Quantification of early artifact, EL-TEP, and VP

We extracted three main features from the response to single TMS pulses applied to the dlPFC. 1) *Early artifact* captured early muscle-related and decay artifacts and was defined as the absolute maximum global mean field amplitude (GMFA) from 12 to 20 ms from the minimally processed data (after epoching and baseline correction) averaged across electrodes, in dBµV [26,33]. 2) Our main outcome measure was the *prefrontal EL-TEP*, defined as the min-to-max amplitude difference within 20 to 60 ms after the TMS pulse and recorded local to the stimulation site (F5, F3, F1, FC5, FC3, FC1; see Figure 1B), in dBµV. We focused on the EL-TEP, which encompasses the P20 and N40 complexes, as a measure of cortical excitability, for several reasons. First, there is intracranial evidence for the neural basis of similar early potentials induced by single pulse electrical stimulation and TMS [44,45]. Second, it has been characterized in the dlPFC [26,27], as well as outside the dlPFC [29,46–48]. Finally, it shows promise in tracking depression pathophysiology and predicting the clinical response to TMS [49,50]. 3) The *vertex potential*, defined as the min-to-max amplitude difference within 80 to 250 ms after the TMS pulse and recorded in the area around the central prefrontal area (FC1, FCz, FC2, C1, C2), allowed evaluation of sensory-related off-target effects [26,31]. Trial-averaged *EL-TEPs*, *early artifact, and vertex potential* were calculated with a 10% trimmed mean approach, excluding trials in the top and bottom 5% of variance to prevent extreme trials from dominating results.

#### 2.6.3 Motor evoked potentials: preprocessing and feature extraction

EMG data recorded during stimulation to M1 were processed using custom scripts in MATLAB 2024b. Epochs were extracted from 900 ms before to 900 ms after each TMS pulse. Data between 1 ms before to 5 ms after each pulse were replaced with values interpolated by autoregressive extrapolation [40] and blending and baseline-corrected based on mean values between 500 to 10 ms before the pulse. Epochs were then high-pass filtered above 1 Hz with a modified filtering approach to reduce the spread of pulse-related artifact into baseline time periods [40] and low-pass filtered below 200 Hz. Line noise was removed by subtraction of a template signal repeating at 60 Hz, with the template constructed from the phase-aligned artifact signal averaged from 50 ms windows shifted at 1/(60 Hz) intervals within each epoch [51]. Peak-to-peak MEP amplitudes were calculated as the minimum-to-maximum voltage from 18 ms to 50 ms post-TMS for each epoch and converted to logarithmic dBmV units. Average MEPs for each intensity were calculated with a 10% trimmed mean.

#### 2.6.4 Source reconstruction

Participant-specific differences in gyral anatomy can cause underlying common cortical sources to project to the scalp in different topographies across participants [52]. To account for this and other related consequences of EEG volume conduction, we performed EEG source estimation. Using digitized electrode locations and individual head models constructed from participants’ anatomical MRI data [37], participant-specific forward models of signal propagation from dipoles distributed over and oriented perpendicular to the cortical surface to electrodes on the scalp were constructed [53,54]. Inverse kernels mapping measured scalp EEG activity to underlying cortical sources were estimated using weighted minimum-norm estimation as implemented in Brainstorm [55]. A source-space region of interest (ROI) capturing the dominant spatial topography of the 45ms EL-TEP response was generated in a data-driven manner based on thresholding to 50% of the max of the group average response magnitude for stimulation at 130% rMT [15]. This ROI was used to calculate source estimates of EL-TEP response magnitudes, with trial averaging performed similarly as described for sensor space data above.

#### 2.6.5 E-field estimation

Electric fields (E-fields) induced by TMS were estimated using SimNIBS [37,38] (v3.2.5). participant-specific head models were generated from T1 and T2 MRIs using headreco, which also generated deformation fields providing a non-linear spatial mapping from MNI coordinates to the brains of individual participants After each session, coil orientations extracted from saved neuronavigation data, combined with a custom dipole model of the TMS coil, were used to predict induced E-fields for each target. Maximum E-field magnitudes on the cortical surface were calculated (after rejecting the top 0.01% of values to remove outliers) to obtain an estimate of stimulation intensity referred to here as |E|. Likewise, maximum E-field components normal (|E_n_|) and tangential (|E_t_|) to the local cortical surface were also calculated.

#### 2.6.6 Input-output curves and sigmoidal curve feature extraction

Following preprocessing and feature extraction (sections 3.1 and 3.2), a Boltzmann sigmoid curve was fit to each participant’s data to characterize the input-output (I/O) curve relating stimulation intensity to cortical responses of the prefrontal or motor cortex. Coefficients were determined using nonlinear least absolute residuals fitting in MATLAB 2023b with the following equation:

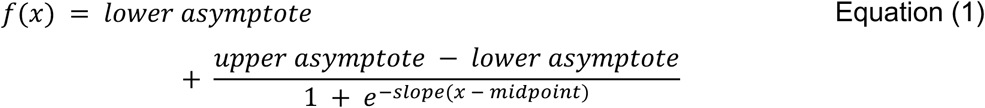

where *f*(*x*) represents the response amplitude and *x* represents the stimulation intensity.

This curve-fitting process generated four parameters: lower asymptote, upper asymptote, midpoint, and slope, along with corresponding 95% confidence intervals for each parameter. The midpoint parameter was selected for subsequent analysis, as it represents the stimulation intensity at which the response is equidistant from its minimum and maximum values, thereby providing a sensitive measure of cortical excitability (Figure 1D). The quality of sigmoidal fitting was evaluated using the midpoint 95% confidence interval (CI) width, with a width <50% rMT considered indicative of sigmoidality.

Additionally, an intensity at half-maximum (IHM) metric was derived from the fitted curves to further characterize cortical excitability without requiring high confidence in sigmoidal fit parameters. The IHM metric was defined as the stimulus intensity eliciting a response amplitude 6 dB below the maximum amplitude (equivalent to a 50% reduction on a linear scale). The maximum amplitude of the fitted sigmoidal curve was first determined within the tested range of stimulus intensities. The half-maximum amplitude was then defined as 6 dB below this maximum value. The IHM was computed as the stimulus intensity at which the fitted curve intersected this half-maximum amplitude, obtained through analytical inversion of the sigmoid function. To ensure physiological validity, two criteria were implemented: (1) the goodness-of-fit must exceed a *R²* threshold of 0.5, ensuring adequate curve fitting, and (2) the calculated IHM must intersect with the fitted curve within the tested intensity range. Participants failing either criterion were excluded from IHM analyses.

### 2.7 Statistical analysis

To characterize the I/O relationship between TMS intensity and EL-TEP amplitude, we employed the sigmoidal curve fitting approach described in section 2.6.6. We report the proportion of participants meeting the sigmoidality criterion (95% confidence interval width <50) and provide descriptive statistics for the midpoint parameter of successful fits. For the I/O curve fitting analysis, data from 24 participants were obtained from their first session, while data from the remaining four participants were obtained from their second session. This selection was necessary to ensure consistency in experimental parameters across all participants: two participants had narrower intensity ranges tested in their initial sessions, and two participants lacked TMS coil angle optimization during their first sessions due to technical issues. These selections were made prior to curve fitting analysis and were based solely on maintaining consistent experimental conditions (60%- 140% intensity range and optimized TMS coil angle) across the cohort.

To compare the effect of different intensities on EL-TEPs, we employed linear mixed-effects models (LME; *N*=28) [56,57], followed by post hoc pairwise comparisons using Bonferroni correction. In our model, intensity level was specified as a fixed effect to assess direct impact, while participants were included as a random intercept term to account for individual differences. This LME approach enabled modeling of within-subject effects with incomplete data, accommodating cases where some intensities were collected only in a subset of participants. We computed estimated marginal means (EMMs) [58] to summarize model values at specific factor levels. Given the unbalanced nature of our data, EMMs represent estimates of the marginal means that would have been observed if the data had been balanced across intensities [58].

To determine whether signal quality influenced EL-TEP I/O curve distributions, we computed signal-to-noise ratio (SNR) as the ratio between EL-TEP amplitude (20-60 ms) and early artifact amplitude (12-20 ms), expressed in decibels (dB). The relationship between signal quality and the ability to detect sigmoidal I/O curves was assessed by comparing sigmoidal and non-sigmoidal groups using the Wilcoxon rank-sum test, a nonparametric method for evaluating distributional differences.

Test-retest reliability analyses were performed on participants who completed two sessions (*n*=13) separated by at least one week. We assessed reliability at both intensity-specific and participant-specific levels by calculating Lin’s concordance correlation coefficient (CCC) [27,59,60], with 95% confidence intervals derived using Fisher’s z-transformation and asymptotic variance estimation as described by Lin [59]. participant-specific CCCs were calculated by comparing first and second visit EL-TEP amplitudes across all available intensity levels for each individual to assess within-participant reliability across different intensities. Intensity-specific reliability was evaluated by calculating CCCs for each stimulation intensity across all participants with paired measurements at that particular intensity level.

To evaluate whether reliable I/O curves could be obtained with fewer trials, we performed systematic subsetting analysis across different trial numbers and intensities. For each intensity level, we retrospectively sampled the first N_t_ trials (N_t_ = 25, 50, 75, 100, 150) from the full set of 200 trials and re-ran preprocessing. Since our extracted response features can systematically vary as a function of number of trials due to variations in noise averaging, we calculated EL-TEP peak-to-peak amplitudes for each of 1000 instances of 25 trials randomly sampled (with replacement) from the N_t_ processed trials (N_t_ = 25, 50,…) and averaged. To assess measurement stability of these subsampled data, we computed CCCs between amplitude values derived from each subset and those from the complete 200-trial dataset. This process was repeated for each intensity level (60-140% rMT). The minimum number of trials needed for reliable measurements was determined by identifying the smallest subset size that produced CCCs > 0.7 with *p* < 0.05 after Bonferroni correction for multiple comparisons across intensities.

To compare parameterizations of I/O curves, we quantified the mean absolute difference (MAD) for each participant between I/O curve response samples and the values predicted by the group-fit I/O curves at matched intensities. For each response measure (sensor-space prefrontal EL-TEP, source-space EL-TEP, and MEP amplitudes), we performed paired *t*-tests between intensity parametrizations (% rMT, % MSO, E-field magnitude, magnitude of E-field normal to the cortical surface, magnitude of E-field tangential to the cortical surface, and % Apex-MT [61]), with Bonferroni correction for multiple comparisons.

To examine relationships between EL-TEPs and other TMS-evoked responses, we performed correlation analyses across EL-TEPs at both sensor and source spaces, MEPs, vertex potentials, and early artifacts. Sigmoidal curve fitting was performed for each of these measures and with stimulation intensity parametrized by % rMT, % MSO, and E-field magnitude. Relationships between IHMs (and midpoints) were assessed using Spearman rank correlations, with separate analyses performed for rMT-normalized amplitude, MSO-normalized amplitude, and E-field magnitude data. Primary correlations of interest included EL-TEP versus MEP relationships, as well as comparisons between EL-TEPs, vertex potentials, and early artifacts.

## 3. Results

### 3.1 Intensity-dependent effects of TMS to the dlPFC

To establish whether prefrontal activation measures exhibit systematic responses to varying TMS intensities, we examined EL-TEPs across multiple intensity levels of TMS to the dlPFC. Individual data revealed a clear, intensity-dependent progression of EL-TEP amplitudes and spatial distributions, with larger responses at higher intensities (Fig. 2A, S1-S2). Both source-level and sensor-level topographies demonstrated progressive increases in response magnitude with increasing stimulation intensity. This pattern was consistently observed at the group level (*N*=28), where topographies and butterfly plots showed similar intensity-dependent effects (Fig. 2B). Examination of EL-TEPs across the full range of stimulation intensities (60-140% rMT) revealed a graded response in both single-subject and group-level data (Fig. 2C, D). The EL-TEP waveforms exhibited distinct peaks within the 20-60 ms time window, with the amplitude of these peaks generally increasing monotonically with TMS intensity. Notably, the group-level data (Fig. 2D) showed a clear separation of EL-TEP amplitudes across intensities, particularly evident for the positive peak around 25 ms and the negative peak around 45 ms post-stimulus. A linear mixed-effects model revealed a significant main effect of TMS intensity on EL-TEP peak-to-peak amplitude (Fig 2E-F, *F*(11,276) = 50.959, *p* < .0001). Post-hoc pairwise comparisons with Bonferroni correction for multiple comparisons showed significant differences between multiple intensity levels (Fig 2F), with the most pronounced differences between lower (60%-90% rMT) and higher intensities (120%-140% rMT; all *p* < 0.001).

**Fig. 2.**
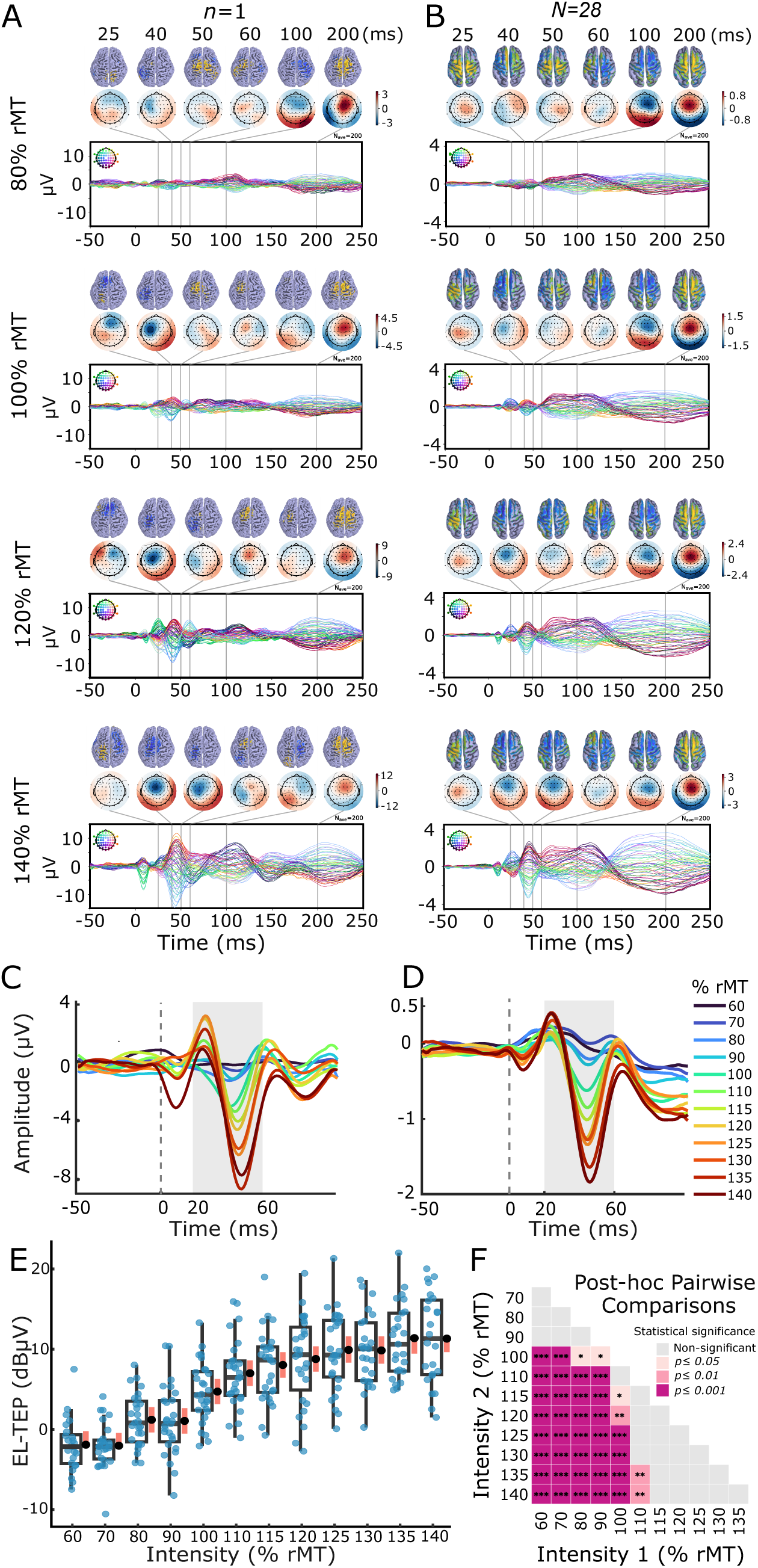
Intensity-dependent effects of TMS to the dlPFC. A) Single-subject source-space and sensor-space topographies and butterfly plots of TMS-evoked potentials (TEPs) at 80%, 100%, 120%, and 140% of resting motor threshold (rMT). Each panel shows the spatial distribution of TEPs in both source space (top) and sensor space (middle) and corresponding butterfly plots with one trace per channel (bottom). B) Group-level (*N*=28) source-level and sensor-level topographies and butterfly plots of TEPs at 80%, 100%, 120%, and 140% rMT. Similar to panel A, but showing the average response across all participants, demonstrating consistent spatial and temporal patterns of activation with increasing stimulation intensity. C) Single-subject EL-TEPs across all stimulation intensities from a left prefrontal sensor-space ROI. Each trace represents the EL-TEP waveform at a different stimulation intensity, illustrating the graded response to increasing TMS intensities. D) Group-averaged EL-TEP waveforms across all stimulation intensities from a left prefrontal sensor-space ROI. Each trace represents the mean EL-TEP waveform at a different stimulation intensity, highlighting the systematic increase in amplitude with higher intensities across the group. Vertical dashed lines in C and D indicate the time of stimulation (0 ms) as well as the time window (20-60 ms) used for EL-TEP amplitude quantification. E) Box plot showing the distribution of EL-TEP amplitudes across stimulation intensities, with individual participant-intensity samples represented in blue. Linear mixed-effects model results showing estimated marginal means (in black) for each stimulation intensity. Orange intervals represent comparison intervals; non-overlapping intervals indicate statistically significant differences (Bonferroni-corrected *p*<0.05) between intensities. F) Matrix visualization of post-hoc pairwise comparisons (Bonferroni-corrected) between stimulation intensities, illustrating significant differences between intensity levels.

### 3.2 Sigmoidal relationship of prefrontal excitability I/O curves

We next characterized the sigmoidality of EL-TEP I/O curves using curve fitting procedures (see Section 2.6.6). Representative individual response curves shown in Fig 3A illustrate the distinct patterns observed across participants: one participant exhibiting a clear sigmoidal I/O curve (teal) with a well-defined threshold and plateau phases, and another displaying a non-sigmoidal response pattern (red) lacking these characteristic features within the sampled range. Individual-level analyses revealed that 57% (16/28) of participants exhibited robust sigmoidal response patterns (Figure 3B left). Critically, the distinction between sigmoidal and non-sigmoidal curves appeared to be related to signal quality, as participants with sigmoidal I/O curves demonstrated significantly enhanced signal-to-noise ratio (SNR, calculated as EL-TEP (in dBµV) minus early artifact (in dBµV)) compared to those with non-sigmoidal responses (Figure 3B right; Wilcoxon rank-sum test, *W* = 143, *p* = .029; *r* = 0.412). Group-level analysis (*N*=28) confirmed sigmoidal relationships across multiple measurement approaches, with group-aggregate sensor-space response curves (Fig 3C-E) exhibiting midpoints at 99% rMT (95% CI: 95-104%), 69% MSO (95% CI: 63-74%), and 114 V/m (95% CI: 100-127 V/m). Similarly, source-space responses as a function of E-field magnitude exhibited a midpoint at 110 V/m (95% CI: 104-116 V/m) (Figure 3F). Additionally, intensity at half-maximum (IHM) values were computed (see Section 2.6.6) to provide a more inclusive characterization of cortical excitability thresholds. While midpoint estimation is limited to participants with well-defined sigmoid curves (CI width <50% rMT), IHM requires only adequate overall curve fit (*R*² > 0.5) and range (>6 dB between min and max), enabling threshold characterization in both sigmoidal and non-sigmoidal participants. IHM values were successfully calculated for 23 of 28 participants (Figure 3C-F). The kernel density estimation plots at the bottom of each panel (Fig 3C-F) illustrate the distribution of midpoints among sigmoidal participants (teal) and IHM values (purple), revealing consistent clustering of neurophysiological thresholds across different measurement approaches. In summary, we found sigmoidal EL-TEP I/O curves in 57% of participants, with sigmoidality predicted by signal quality and group-level curves exhibiting midpoints near motor threshold.

**Fig. 3.**
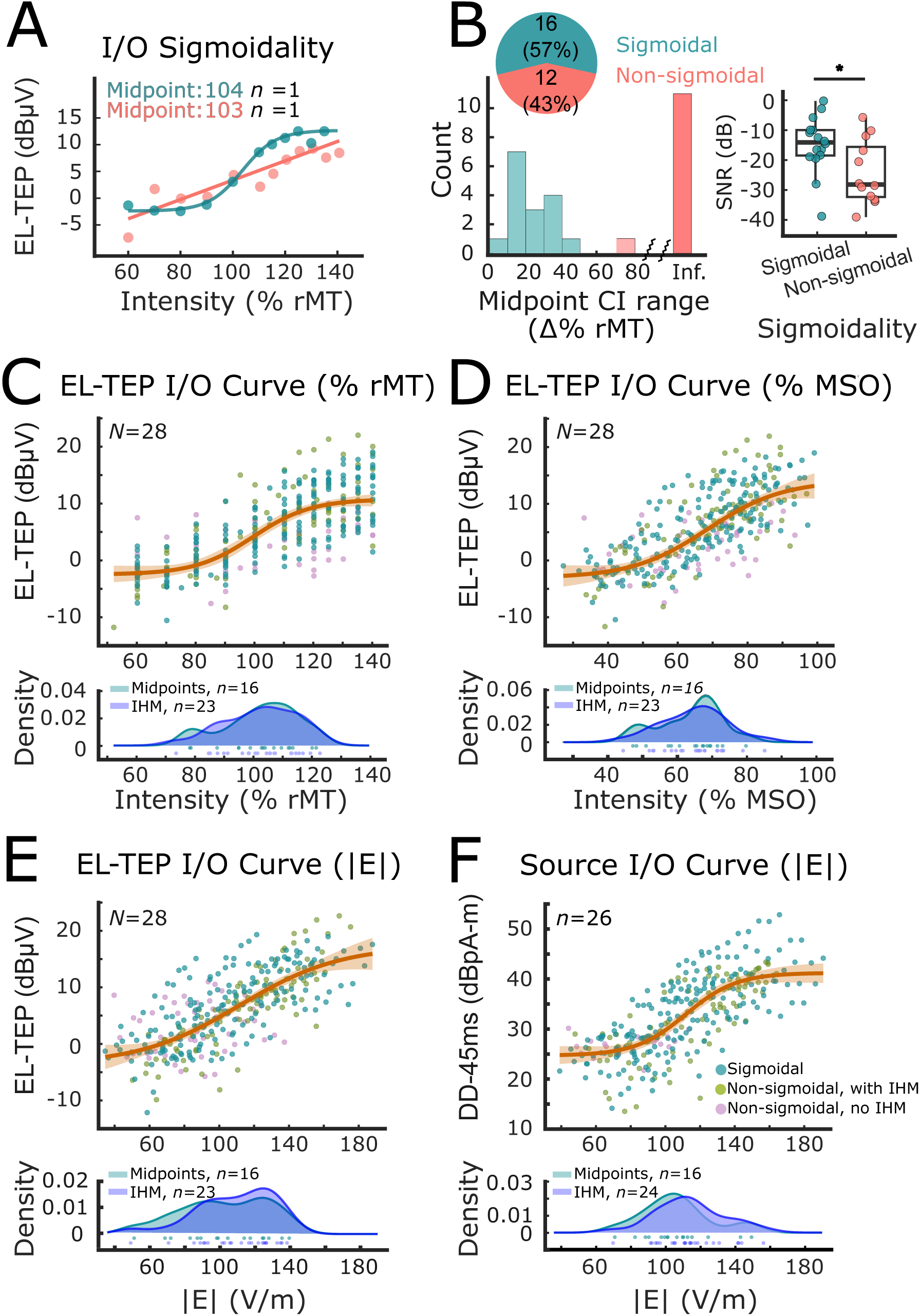
Sigmoidal relationship of prefrontal excitability I/O curves. (A) Representative individual I/O curves showing sigmoidal (teal) and non-sigmoidal (red) examples, *n*=1 participant each. (B) Top-left: pie chart showing the proportion of sigmoidal (57%, *n*=16) and non-sigmoidal (43%, *n*=12) participants, with the histogram below displaying the distribution of 95% confidence interval ranges for midpoint estimation. Right: box plot showing significantly higher signal-to-noise ratios (SNR; EL-TEP minus artifact, in dB) in sigmoidal participants compared to non-sigmoidal participants. (C-F) Group-level input-output relationships between TMS intensity and EL-TEP amplitude (*N*=28), showing sigmoidal curves with teal dots representing sigmoidal participants, green dots representing non-sigmoidal participants with valid IHMs, and pink dots representing non-sigmoidal participants without IHMs. Kernel density plots at the bottom of each panel illustrate the distribution of midpoints for sigmoidal participants (teal) and IHM values (purple). (C) EL-TEP amplitude versus TMS intensity (% rMT) showing a sigmoidal curve with the midpoint at 99% rMT (95% CI: 95-104%). (D) EL-TEP amplitude versus absolute TMS intensity (% MSO) showing a sigmoidal curve with the midpoint at 69% MSO (95% CI: 63-74%). (E) EL-TEP amplitude versus electric field magnitude (|E| V/m) showing a sigmoidal relationship with the midpoint at 114 V/m (95% CI: 100-127 V/m). (F) Source-level responses (Data driven at 45ms, dBμA-m) versus electric field magnitude (|E| V/m) with the midpoint at 110 V/m (95% CI: 103-116 V/m) (*n*=26). * indicates *p* < 0.05.

### 3.3 Prefrontal EL-TEPs show greater reliability at higher stimulation intensities

To establish the stability of prefrontal EL-TEPs across sessions, we conducted test-retest reliability analyses across at least two weeks in a subset of participants (*n*=13, see Section 2.7). Examples of I/O curves acquired from one individual on separate days are shown in Fig 4A, with the correlation between responses at matched intensities shown in Fig 4B. Additional results are shown in Fig S3-S4. Within-subject test-retest reliability measures, calculated based on all available matched intensities, showed considerable variation across participants, with CCCs ranging from -0.23 to 0.94 (Fig. 4C), with a median CCC of 0.77. Single-intensity across-subject reliability at 140% rMT demonstrated strong correlation between session measurements (CCC = 0.723, *n*=12; Fig. 4D). Analysis of reliability across the full range of intensities revealed systematically improved test-retest reliability at higher stimulation intensities, with consistently high reliability (CCC > 0.5) achieved at intensities above 100% rMT while lower intensities showed substantially weaker reproducibility (Fig. 4E). In summary, the strong test-retest reliability across weeks supports EL-TEPs as a robust marker of prefrontal excitability, especially when measured with higher intensity TMS.

**Fig. 4.**
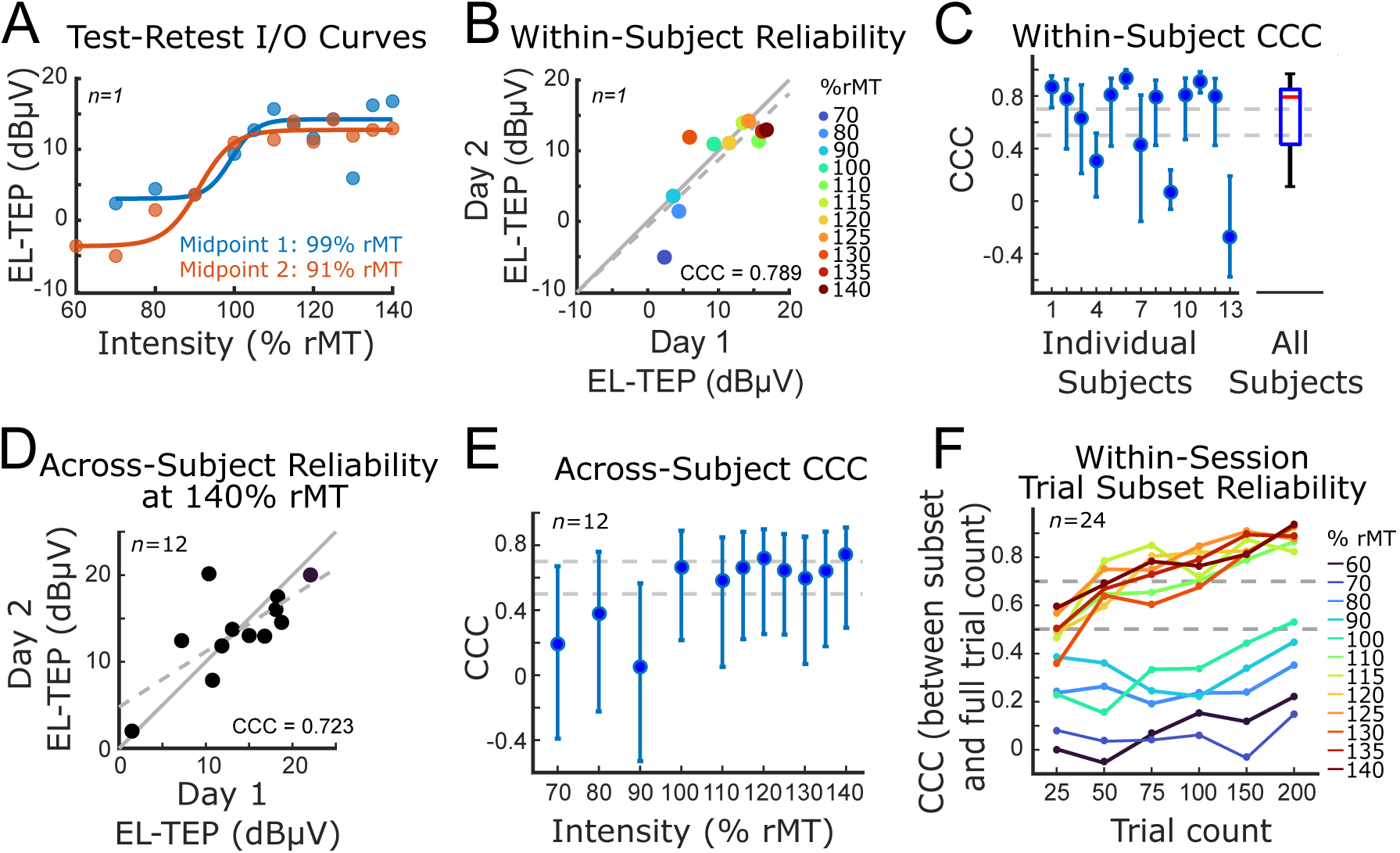
Prefrontal excitability I/O curves are reliable and obtainable with a subset of trials and intensities. (A) Representative input-output (I/O) curves from a single participant showing EL-TEP amplitudes plotted against stimulation intensity (% rMT) for two separate sessions (blue and orange). Sigmoid curve fits demonstrate consistent response profiles with similar midpoints. (B) Within-subject reliability across intensities for a representative participant showing strong correlation between first and second session EL-TEP measurements. Colored dots represent different stimulation intensities ranging from 70-140% rMT. (C) Left: Individual concordance correlation coefficients (CCC) for test-retest reliability across 13 participants, with bars indicating 95% confidence intervals. Right: Box plot summarizing within-subject reliability across all participants (median CCC = 0.77, 95% CI = [0.395, 0.830]). (D) Across-subject reliability at 140% rMT (*n*=12) demonstrating strong correlation between session 1 and session 2 measurements. (E) Intensity-dependent reliability across subjects (*n*=12) showing systematically improved test-retest reliability at higher stimulation intensities, with consistently high reliability (CCC > 0.5) achieved at intensities of 100% rMT and higher. Bars indicate 95% confidence intervals. (F) Within-session trial subset reliability comparing different trial subsets to the full trial count (*n*=24). Each line represents reliability at a specific stimulation intensity (60-140% rMT). At higher intensities (>100% rMT), as few as 50 trials achieved reliability of CCC>0.5.

To determine whether reliable responses could be obtained with fewer TMS pulses (trials), we next evaluated the relationship between trial count and measurement stability. Within- session analysis of trial subsets (*n*=24) revealed that as few as 100 trials per intensity could provide reliable EL-TEP measurements, particularly at higher stimulation intensities (Fig. 4F). The stability of measurements showed a clear intensity-dependent pattern, with higher intensities demonstrating robust reliability even with reduced trial counts. Specifically, at stimulation intensities below 110% rMT, correlations between trial subsets (25, 50, 75, 100, 150 trials) and the full trial sets (200 trials) were minimal (CCC < 0.5). In contrast, stimulation intensities at 110% rMT and higher demonstrated more robust reliability (CCC > 0.5) with at least 50 trials. Together, these results suggest that protocols using fewer trials at higher intensities could still provide reliable characterization of prefrontal excitability.

### 3.4 Alternate parameterizations of I/O curves

To determine which parameterizations of stimulation intensity best capture inter-individual variability in I/O curves across response measures, we compared multiple intensity parameterizations across sensor-space EL-TEPs, source-space EL-TEPs, and MEPs. Stimulation intensity can be quantified in multiple ways, including relative to rMT and MSO and in terms of estimated maximum E-field magnitude at the cortical surface. TMS-evoked responses can be quantified in terms of MEP amplitude (when stimulating motor cortex), or in terms of EEG scalp or source-space TEP responses at various latencies. Figure 5 (A-C,E-G,I-K) shows group average and individual I/O curves for several combinations of these input and output parameterizations. These plots provide a visualization of population variability in I/O curves relative to the group means for each parameterization. We quantified the amount of variability in responses not explained by the fitted group-level I/O curves with the mean absolute difference (MAD) between response samples for each participant and predictions at corresponding intensities along the group I/O curve, as shown in Figure 5 (D,H,L). There were no significant MAD differences between parameterizations for sensor-space (Fig 5 A-D) or source-space (Fig 5 E-H) EEG responses. In considering EMG responses (Fig 5 I-L), % rMT was the only significantly different parameterization, with lower MADs than all others (mean differences in MAD of 3.39, 6.86, 6.71, and 6.84 dB and Bonferroni-adjusted p values of 0.0248, 0.000119, 0.000237, and 0.000136 when compared to % MSO, |E|, |En|, and |Et| respectively); this was expected, since the % rMT values were directly normalized by measurement of participant-specific motor thresholds at the beginning of sessions. The next largest differences were between % MSO parameterization and E-field derived parameterizations of MEP responses, but these differences were not significant (mean differences in MAD of 3.465, 3.33, and 3.46 dB and Bonferroni-adjusted *p* values of 0.0979, 0.118, and 0.0882 when compared to |E|, |En|, and |Et| respectively). In summary, intensity parameterizations showed no significant differences for EEG responses, while % rMT was superior for MEP responses as expected.

**Figure 5.**
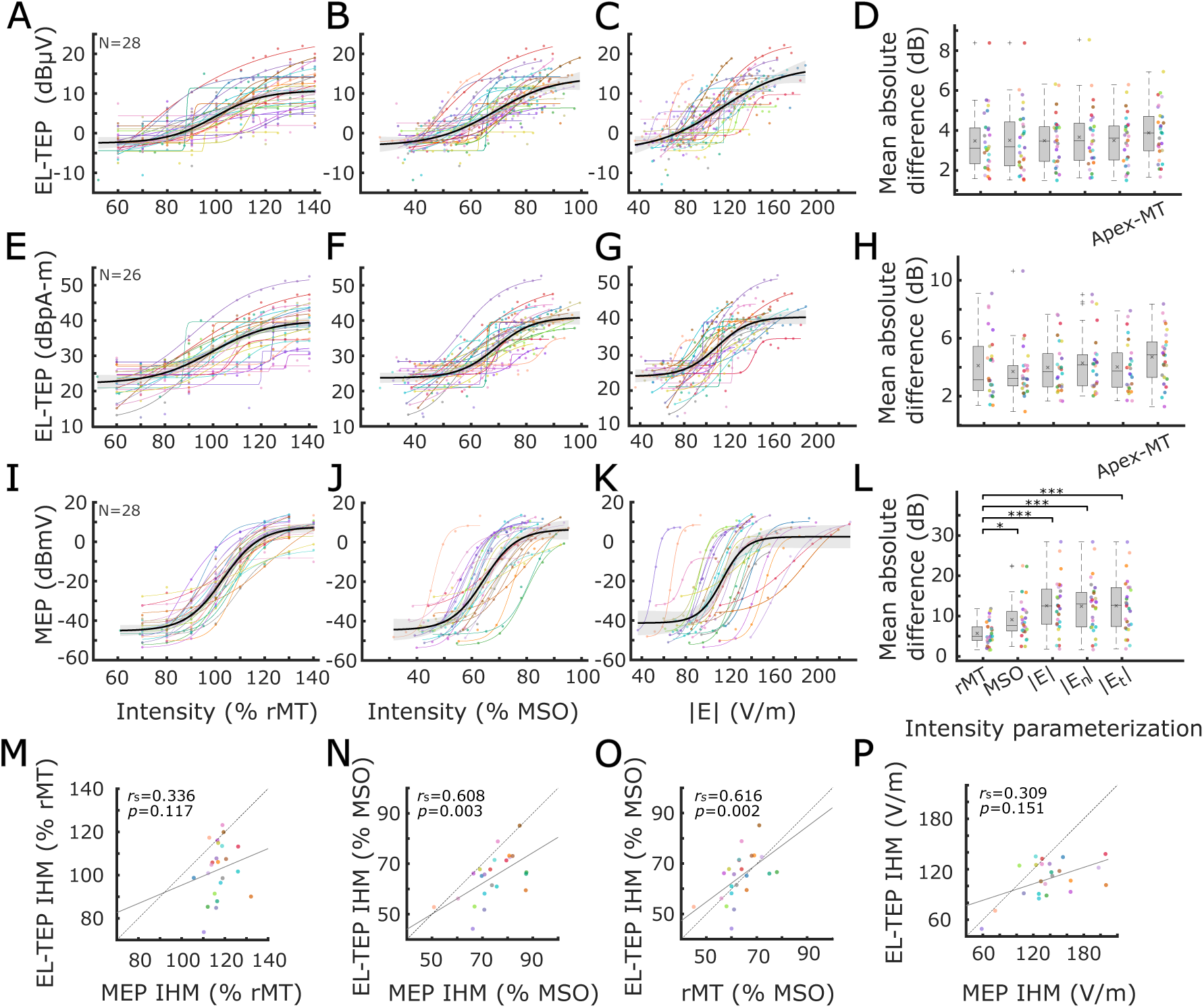
Alternate parameterizations of I/O curves and relationships between EL-TEPs and MEPs. Input-output curves depicting sensor-level EL-TEP responses across intensity metrics: % rMT (A), % MSO (B), and E-field magnitude (C). Input-output curves for source-level EL-TEP responses across identical intensity metrics (E, F, G). Input-output curves for MEP responses across identical intensity metrics (I, J, K). Colored lines represent individual participants’ fitted sigmoidal curves, with the black line denoting the group-averaged response. Mean absolute differences between individual responses and group mean I/O curves are quantified for sensor-level EL-TEP responses (D), source-level EL-TEP responses (H), and for MEP responses (L), with scatter plot colors matching colors in the I/O curves to the left. Each middle horizontal line in the boxplots represents the median, and X represents the mean of MAD values. Asterisks represent Bonferroni-adjusted *p* values of *p*<0.05 (*) and *p*<0.001 (***). Scatter plots (M-P) illustrate correlations between sensor-level EL-TEP curve intensity at half-maximum (IHM) values MEPs, with MEPs parameterized by % rMT IHM (M, Spearman rho = 0.336, *p* = 0.117), % MSO IHM (K, Spearman rho = 0.608, *p* = 0.003), estimated resting motor threshold (rMT, O), and E-field IHM (P, Spearman rho = 0.309, *p* = 0.151). Each data point represents an individual participant, with the dotted diagonal line representing identity.

### 3.5 Relationship between EL-TEPs and other TMS-evoked responses

To better understand how EL-TEPs relate to other measures of cortical response to TMS, we examined their relationship with early artifacts (AF) and vertex potentials (VP) across different intensity metrics (Fig. S5). Plots of individual I/O curves across all response measures are included in Fig S6-S7. Analysis of group-level I/O curves revealed distinct patterns across these measures when examined against % rMT, % MSO, and E-field strength. While EL-TEPs demonstrated characteristic sigmoidal relationships with intensity, AF and VPs displayed different I/O curve characteristics. 54% (15/28) of participants exhibited sigmoidal response patterns of AF (Figure S5A). Only 25% (7/28) of participants exhibited sigmoidal response patterns of VP (Figure S5E), showing more variable responses with often linear patterns.

We quantified the differences between participant-specific AF and VP I/O curves and group-averages with MAD (Fig S5D,H), as described above for EL-TEP and MEP curves (see sections 2.7, 3.4). After Bonferroni correction for multiple comparisons, the only significant difference in AF MADs was between % rMT and % Apex-MT parameterizations (MAD difference 1.15 dB, adjusted *p*=0.0397); all other adjusted *p* values were > 0.1.

When examining correlations between EL-TEP and AF IHMs, we observed significant relationships for % MSO (Fig. S5J; Spearman rho = 0.583, *p* = 0.004) and E-field strength (Fig. S5K; Spearman rho = 0.850, *p* < 0.001), while the correlation with % rMT was not significant (Fig. S5I; Spearman rho = 0.291, *p* = 0.178). Between EL-TEP and VP IHMs, we found significant correlations for % MSO (Fig. S5M; Spearman rho = 0.519, *p* = 0.021) and E-field strength (Fig. S5N; Spearman rho = 0.815, *p* < 0.001), while % rMT showed no significant correlation (Fig. S5L; Spearman rho = 0.265, *p* = 0.258). Together, these findings suggest that when representing stimulation intensity on an absolute scale (% MSO or E-field magnitude), participants with larger EL-TEP IHMs also tended to have larger AF and VP IHMs, and this correlation was not apparent after normalizing intensity by motor threshold.

## 4. Discussion

### 4.1 Summary of findings

This study provides the first systematic characterization of prefrontal excitability input-output relationships in humans using early local TMS-evoked potentials (EL-TEPs). Our data support that dlPFC EL-TEPs are stimulation intensity dependent (Fig 2) and that I/O curves are sigmoidal in the majority of participants. Additionally, we found that detection of sigmoidal I/O curves was strongly predicted by signal quality (Fig 3) We found variable within-subject test-retest reliability of EL-TEPs across all intensities, but strong reliability at higher stimulation intensities with as few as 50-75 trials per intensity (Fig 4) EL-TEP curves shared some properties with motor-evoked potentials, but parametrization of I/O curves using % rMT did not significantly reduce inter-individual variability compared to non-motor parametrizations such as % MSO or E-field magnitude (Fig 5). EL-TEPs showed distinct curves from early artifacts and vertex potentials.

### 4.2 Sigmoidal I/O curves of the EL-TEP

The sigmoidal relationships we observed between TMS intensity and EL-TEP amplitude suggest that the prefrontal cortex follows nonlinear recruitment patterns similar to those seen in other cortical regions [19,21,22]. Critically, the detection of these sigmoidal relationships was strongly dependent on signal quality – participants with high signal-to-noise ratios consistently demonstrated EL-TEP amplitudes that increased nonlinearly with TMS intensity, exhibiting distinct phases of minimal response, rapid growth, and saturation that mirror the canonical input-output properties of neuronal populations [21,62,63]. This interpretation is supported by converging evidence from other methodologies – previous studies using intracranial recordings have shown that direct electrical stimulation of human cortex produces similar sigmoidal recruitment patterns [64,65], while computational models suggest these curves reflect the interaction between excitatory and inhibitory circuits within local cortical regions [66,67]. This framework could be particularly valuable for understanding altered prefrontal excitability in psychiatric conditions and for titrating therapeutic interventions to individual physiological parameters rather than relying on standardized protocols.

### 4.3 Reliability and optimization of EL-TEP I/O curves

Our reliability analyses revealed important insights into the measurement properties of EL-TEP I/O curves. Within-subject across-intensity reliability showed considerable variability, indicating that individual participants’ response profiles varied in their consistency across different stimulation intensities. Across-subject per-intensity reliability demonstrated a clear intensity-dependent pattern, with consistently better reliability at stimulation intensities above resting motor threshold. While previous studies have established the reliability of single-intensity TEP measurements in motor and prefrontal regions [60,68], our findings extend this work by demonstrating that reliability varies systematically across the intensity spectrum, with higher intensities in general providing more stable measurements.

Building on this intensity-dependent reliability pattern, trial subset analysis revealed that reliable EL-TEP responses can be obtained with as few as 50-75 TMS pulses at high intensities (≥110% rMT). This represents a substantial improvement in efficiency compared to previous TEP studies, which have typically used 150-200 pulses for single-intensity measurements [26,27,31,33]. Specifically, at stimulation intensities below 110% rMT, correlations between subsets and full trial sets were minimal (CCC < 0.5), whereas stimulation intensities at 110% rMT and higher demonstrated robust reliability (CCC > 0.5 when using ≥ 50 trials). These findings can inform the choice of stimulation intensity and number of pulses needed for reliable TMS-EEG assessments, with important implications for both research and clinical applications, especially for rapid characterization of individual prefrontal I/O relationships and titrating therapeutic interventions to individual physiological responses rather than relying on standardized protocols.

### 4.4 Relationship between EL-TEPs and corticospinal excitability

The relationship between prefrontal and corticospinal excitability has important implications for optimizing noninvasive brain stimulation protocols. A key finding of our study was a strong positive correlation between MEP and EL-TEP intensity at half-maximum (IHM) values in % MSO (Fig 5N; Spearman rho = 0.608, p = 0.003). This correlation between prefrontal and motor excitability thresholds likely reflects multiple interacting mechanisms. The robust % MSO correlation, combined with weaker correlations when intensities were quantified by electric field magnitude (Fig 5P; Spearman rho = 0.309, p = 0.151), suggests that anatomical factors such as skull thickness and coil-to-cortex distance may have contributed significantly to individual differences in stimulation thresholds. However, this does not preclude additional contributions from shared neurophysiological properties between motor and prefrontal cortices, including common cellular architecture, inhibitory circuits, and cortical excitability mechanisms [69–72].

Importantly, while both TEP and MEP measures exhibited sigmoidal relationships with increasing stimulation intensity, EL-TEP curves showed distinct characteristics including shallower slopes compared to MEP curves, indicating region-specific aspects of excitability that complement the shared threshold properties. Our analysis of different intensity parameterizations revealed that no single approach significantly better accounted for inter-individual variability in EL-TEP responses (Fig 5D,H), highlighting the complex, multifactorial nature of this cortical excitability measure. For MEPs, only parameterizing intensity by % rMT (itself already a measure of an I/O curve threshold) produced significantly lower inter-individual variability than other parameterizations. Critically, personalized computational modeling to predict induced electric fields were no better than % MSO in explaining inter-individual variability of responses (Fig 5L).

These findings have important clinical implications. The non-superiority of any one intensity parametrization in describing inter-individual EL-TEP response variability motivates a need to improve computational models to better account for stimulation effects on the brain [73–77], for mapping the direct cortical and subcortical effects of TMS [78], and understanding how these effects map to signals measured with EEG scalp electrodes [79–82]. These results point toward the value of direct physiological measurement of prefrontal responses rather than relying solely on anatomical proxies or motor-based estimates.

### 4.5 EL-TEPs capture neural responses distinct from sensory and artifactual effects

A critical challenge in measuring prefrontal excitability with TMS-EEG is distinguishing true neural responses from non-neural artifacts and sensory effects. Our analyses demonstrated that EL-TEPs represent distinct physiological responses that are dissociable from both early artifacts and later sensory effects. In addition to their previously-reported spatial and temporal differences, we showed distinct input-output relationships between EL-TEPs, early artifacts, and vertex potentials quantified by IHM thresholds (Fig S5). Collectively, the distinct response properties of EL-TEPs compared to both artifactual and sensory signals suggest that EL-TEPs capture a unique measure of local cortical excitability, providing a foundation for their use as physiological biomarkers in both research and clinical applications.

### 4.6 Clinical implications for monitoring and optimizing prefrontal interventions

The establishment of reliable prefrontal I/O curves has significant implications for improving neuromodulatory treatments of psychiatric disorders. Specifically, our finding that individual prefrontal excitability may be efficiently characterized through EL-TEP I/O curves opens new possibilities for personalized medicine approaches to TMS therapy. Current standardized treatment protocols rely on motor threshold for intensity selection but do not account for individual variations in prefrontal physiology that may influence treatment response. The ability to directly measure prefrontal I/O relationships could enable more precise titration of stimulation parameters for individual patients. Additionally, our finding that reliable EL-TEP responses can be obtained with as few as 50-75 pulses per intensity level makes it feasible to incorporate these measurements into clinical workflows, potentially enabling monitoring of these prefrontal excitability measures during and after treatment. These advances could help address the significant variability in treatment outcomes currently observed in TMS therapy for depression [83,84].

### 4.7 Limitations and future directions

Several important limitations of the current work need to be considered. First, our findings are based on a relatively small sample of healthy participants, and the generalizability to psychiatric populations remains to be established. Second, the significant variability in signal quality across participants, with only a narrow majority showing clear sigmoidal relationships, highlights the ongoing challenge of obtaining robust TMS-EEG measurements in individual participants. Third, while our sample size was adequate for demonstrating the feasibility of I/O curve measurements, larger studies will be needed to establish normative ranges and reliable clinical thresholds. Fourth, our study was limited to a single prefrontal target (dlPFC), and it remains unclear whether similar I/O curve properties would be observed across other prefrontal subregions or cortical areas. Fifth, our measurements were obtained during one or two sessions per participant under controlled laboratory conditions, and the stability of these curves across different brain states, times of day, and clinical contexts requires further investigation.

Looking forward, this work opens several promising avenues for future investigation. First, systematic mapping of EL-TEP I/O curves across different dlPFC subregions could help optimize stimulation target selection, particularly given the known variability in treatment response across different prefrontal targets [85–87]. Second, investigating how these curves vary with brain state and arousal could provide insights into optimal treatment timing and conditions. Third, characterizing alterations in prefrontal I/O curves across psychiatric disorders could yield valuable biomarkers for patient stratification and treatment selection. Fourth, longitudinal studies examining how these curves change during successful treatment could enhance our understanding of therapeutic mechanisms and enable real-time treatment optimization.

Finally, a particularly promising avenue for advancing our understanding of prefrontal dose-response mechanisms lies in intracranial investigations combining simultaneous dlPFC TMS with intracranial EEG and single-unit recordings. Such approaches could provide unprecedented insight into the cellular and circuit-level mechanisms underlying the sigmoidal relationships we observed with scalp EEG. Recent advances in combining TMS with intracranial recordings have begun to reveal the neural basis of cortical excitability measures [78,88,89], while studies using direct electrical stimulation with intracranial recordings have demonstrated the feasibility of characterizing dose-response relationships [90]. Applying similar dose-response curve methodologies to these intracranial approaches could bridge the gap between our scalp-based EL-TEP measurements and the underlying neurophysiological mechanisms. This mechanistic understanding could ultimately inform more precise targeting strategies and improve our ability to predict individual treatment responses based on underlying circuit properties.

## 5. Conclusions

This study provides the first systematic characterization of prefrontal I/O relationships using EL-TEPs, demonstrating clear sigmoidal input-output properties in over half of participants (57%), with signal quality strongly predicting curve detection. While we observed correlations between motor and prefrontal excitability measures in aggregate, inter-individual variability in the relationship between motor and prefrontal excitability was high. Comparison across intensity parameterizations motivates the need for individualized monitoring of stimulation responses. Reliable prefrontal excitability measurements could be obtained efficiently with as few as 50-75 pulses per intensity level, making it feasible to quickly collect EL-TEP I/O curves for assessing prefrontal cortex function. This framework enables a shift from motor-proxy-based stimulation protocols toward direct measurement of target region responses, opening new possibilities for more precise and personalized brain stimulation treatments for psychiatric disorders.

## Acknowledgements.

We extend gratitude to all of our research participants. We would also like to acknowledge the generous contributions of the members of the Personalized Neurotherapeutics Laboratory for helpful feedback on the manuscript and throughout the course of the study. This research was supported by the National Institute of Mental Health under award number R01MH126639, R01MH139650, R01MH129018, and a Burroughs Wellcome Fund Career Award for Medical Scientists (CJK). NFC was supported by the Stanford Taiwan Science and Technology Hub Fellowship. JMR was supported by the Department of Veterans Affairs Office of Academic Affiliations Advanced Fellowship Program in Mental Illness Research and Treatment, the Medical Research Service of the Veterans Affairs Palo Alto Health Care System and the Department of Veterans Affairs Sierra-Pacific Data Science Fellowship. JG was supported by personal grants from Orion Research Foundation, the Finnish Medical Foundation, and Emil Aaltonen Foundation.

## SUPPLEMENTARY INFORMATION

### Table S1 Demographics

**S1.1.**
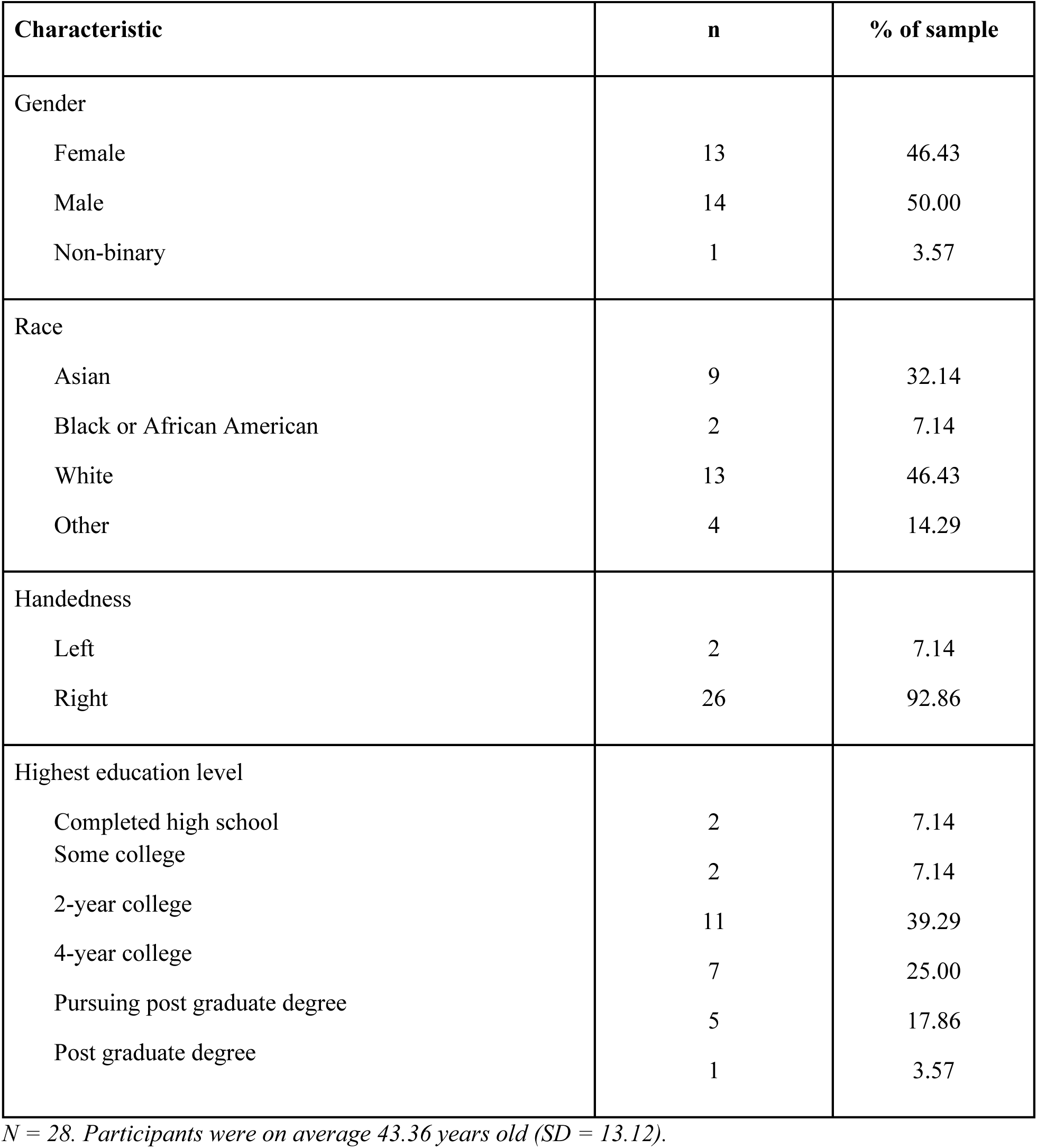
Full sample, *N=28*.

**S1.2.** Reliability testing sample, *n=13*.

**S1.2 Reliability testing sample, $n=13$**
| Characteristic | n | % of sample |
| --- | --- | --- |
| Gender |  |  |
| Female | 5 | 38.46 |
| Male | 8 | 61.54 |
| Race |  |  |
| Asian | 3 | 23.08 |
| Black or African American | 2 | 15.38 |
| White | 7 | 53.85 |
| Other | 1 | 7.69 |
| Handedness |  |  |
| Right | 13 | 100.00 |
| Highest education level |  |  |
| Completed high school | 1 | 7.69 |
| 4-year college | 5 | 38.46 |
| Pursuing post graduate degree | 2 | 15.38 |
| Post graduate degree | 5 | 38.46 |

$n = 13$ . Participants were on average 46.69 years old ( $SD = 12.11$ ).

---

**Fig. S1.**
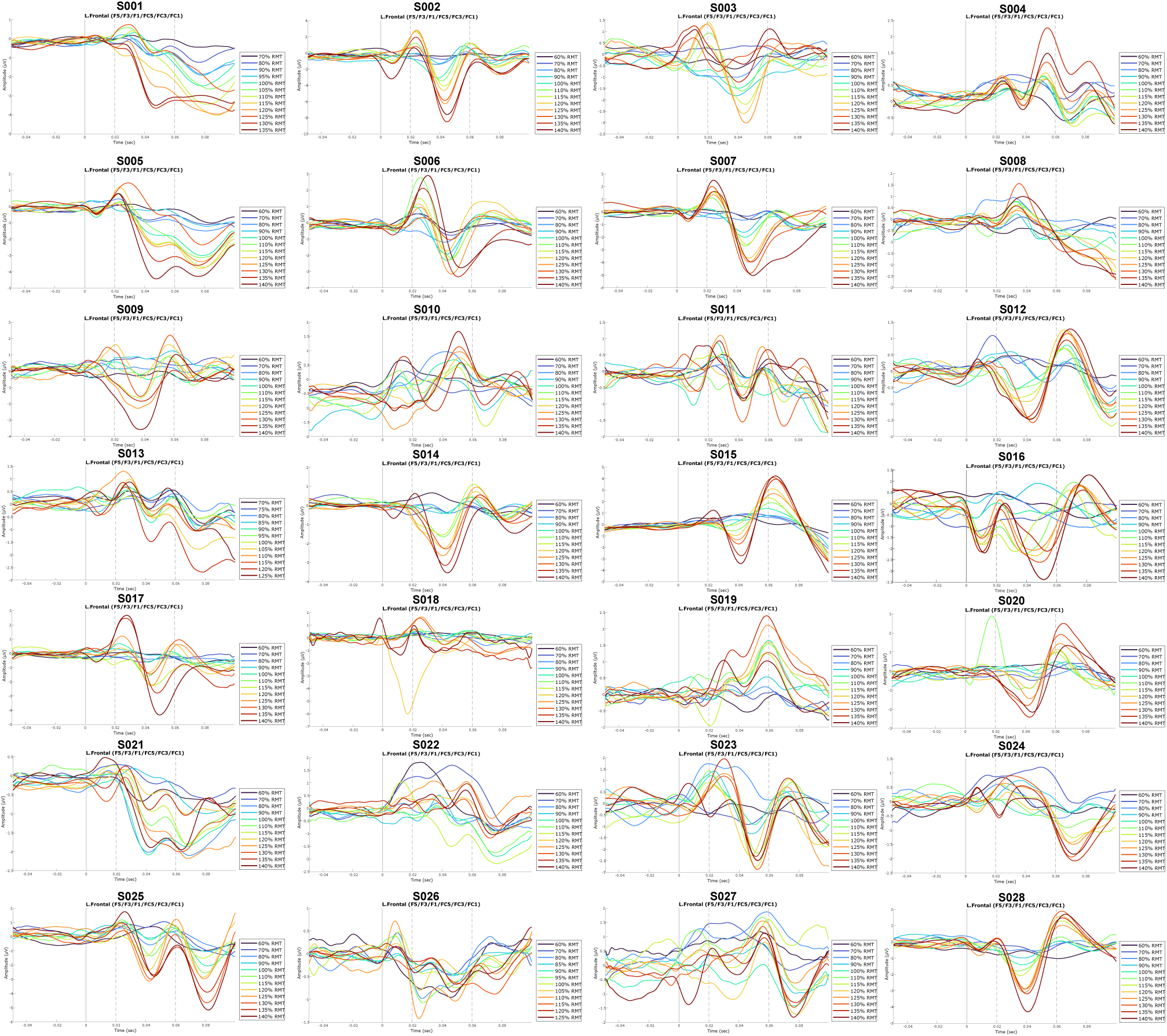
Individual EL-TEP waveforms. Early local TMS-evoked potentials (EL-TEPs) recorded from individual participants, showing the characteristic positive peak around 25ms and negative peak around 45ms across different stimulation intensities (60-140% resting motor threshold).

**Fig. S2.**
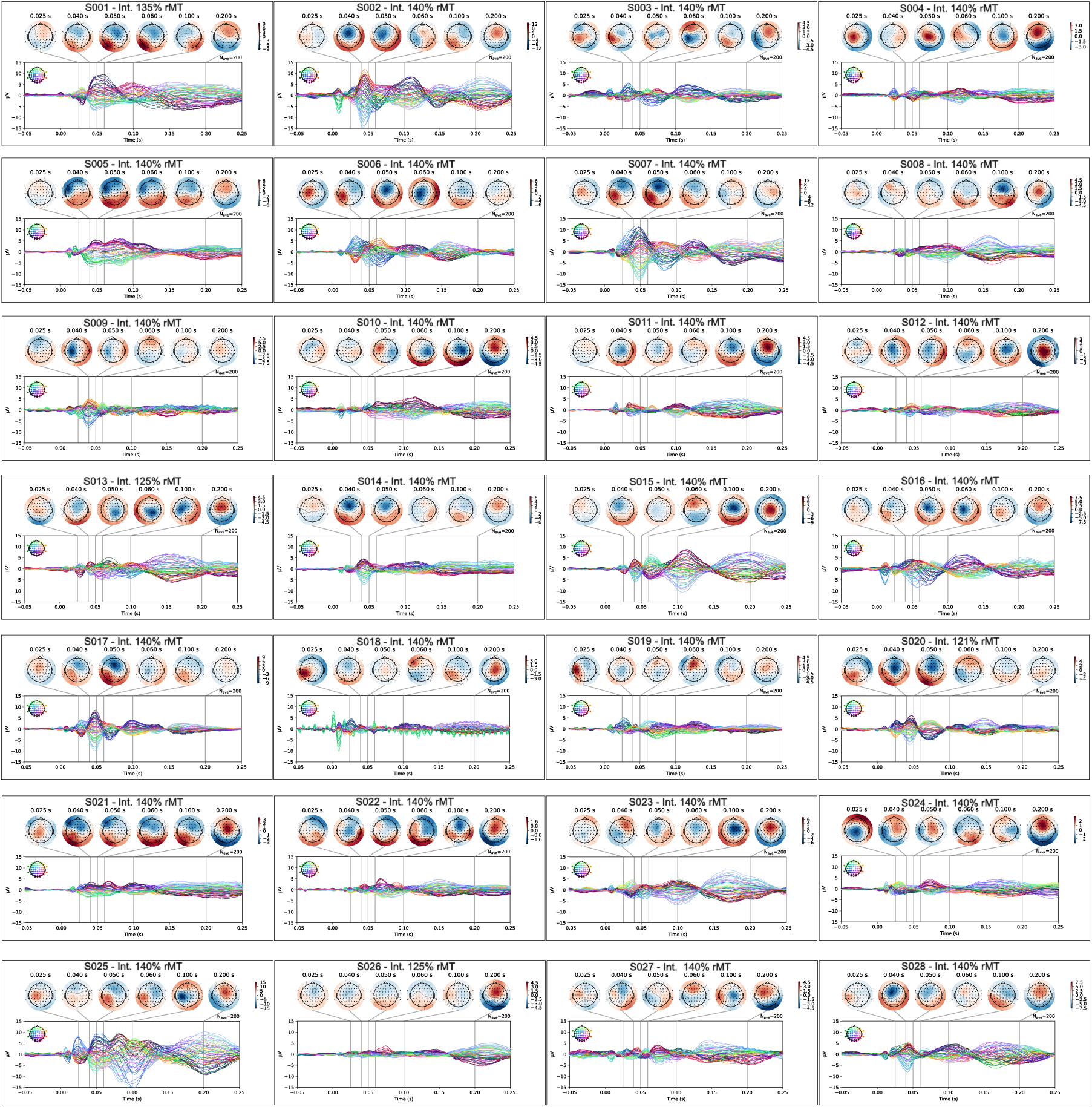
Individual Time Topography at maximum intensity. Topographical plots demonstrating the spatial distribution of TMS-evoked responses at 130% resting motor threshold for each participant, illustrating the consistency of localized activation patterns in the stimulated prefrontal region.

**Fig. S3.**
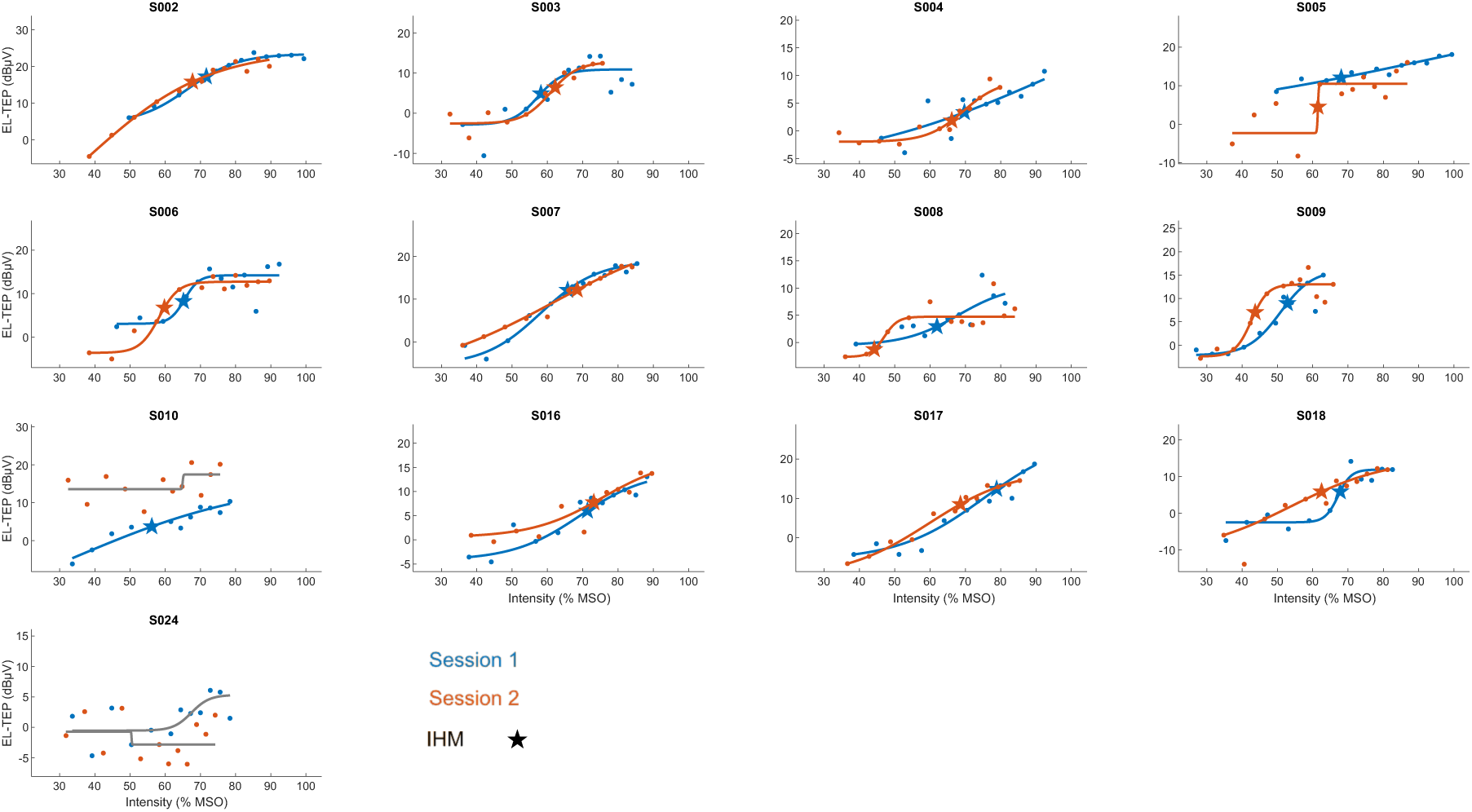
Individual test-retest I/O curves for EL-TEP across sessions. Individual input- output curves from 13 participants showing EL-TEP amplitude as a function of TMS intensity (% MSO) across two sessions separated by at least two weeks. Blue dots and lines represent session 1, orange dots and lines represent session 2. Stars indicate intensity at half-maximum (IHM) values for each session when calculable.

**Fig. S4.**
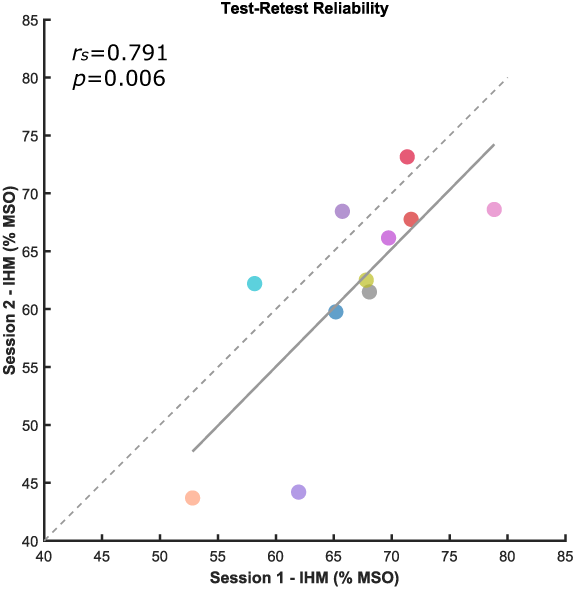
Test-retest reliability of intensity at half-maximum (IHM) values. Scatterplot showing correspondence between session 1 and session 2 IHM values (expressed as % MSO) for 13 participants. Each colored dot represents one participant. The gray dashed line represents perfect agreement (identity line), while the solid gray line shows the linear regression fit. Strong test-retest reliability was observed (Spearman rs = 0.791, p = 0.006), indicating that IHM provides a stable metric of prefrontal excitability threshold across sessions.

**Fig. S5.**
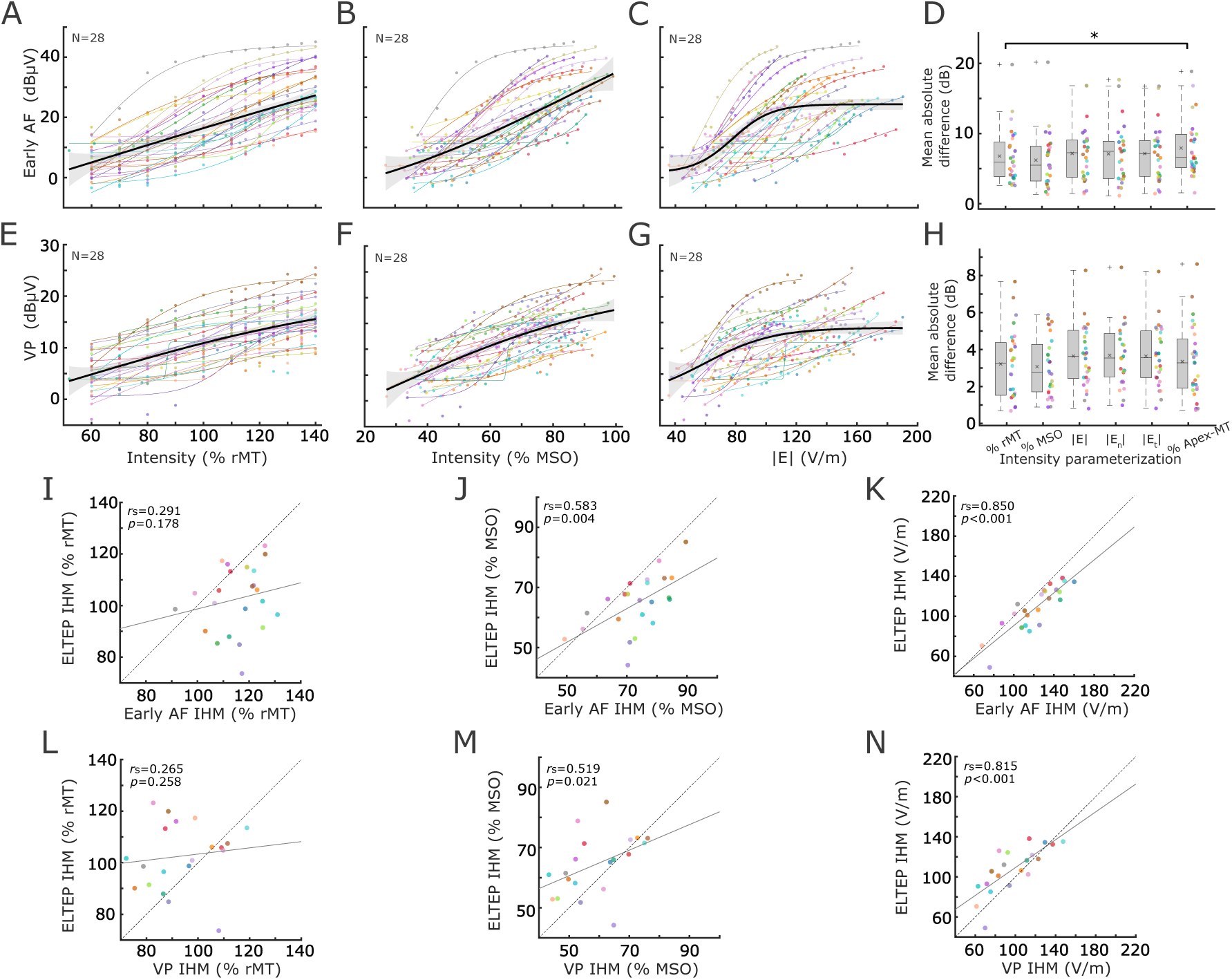
Alternate parameterizations of I/O curves and relationships between EL- TEPs and early artifact, vertex potentials. Group-level input-output curves for early artifacts (AF) across intensity parameterizations: % rMT (A), % MSO (B), and E-field magnitude (C), with box plots (D) showing mean absolute differences between individual responses and group mean I/O curves for AF responses. Group-level input-output curves for vertex potentials (VP) across intensity parameterizations: % rMT (E), % MSO (F), and E-field magnitude (G), with box plots (H) showing mean absolute differences between individual responses and group mean I/O curves for VP responses. Scatter plots illustrating correlations between EL-TEP and early artifact intensity at half-maximum (IHM) values for % rMT (I), % MSO (J), and E-field magnitude (K). Scatter plots showing correlations between EL-TEP and vertex potential IHM values for % rMT (L), % MSO (M), and E-field magnitude (N). Colored lines in I/O curves represent individual participants’ fitted sigmoidal curves, with black lines denoting group-averaged responses. Correlation coefficients and p-values are displayed on each scatter plot panel.

**Fig. S6.**
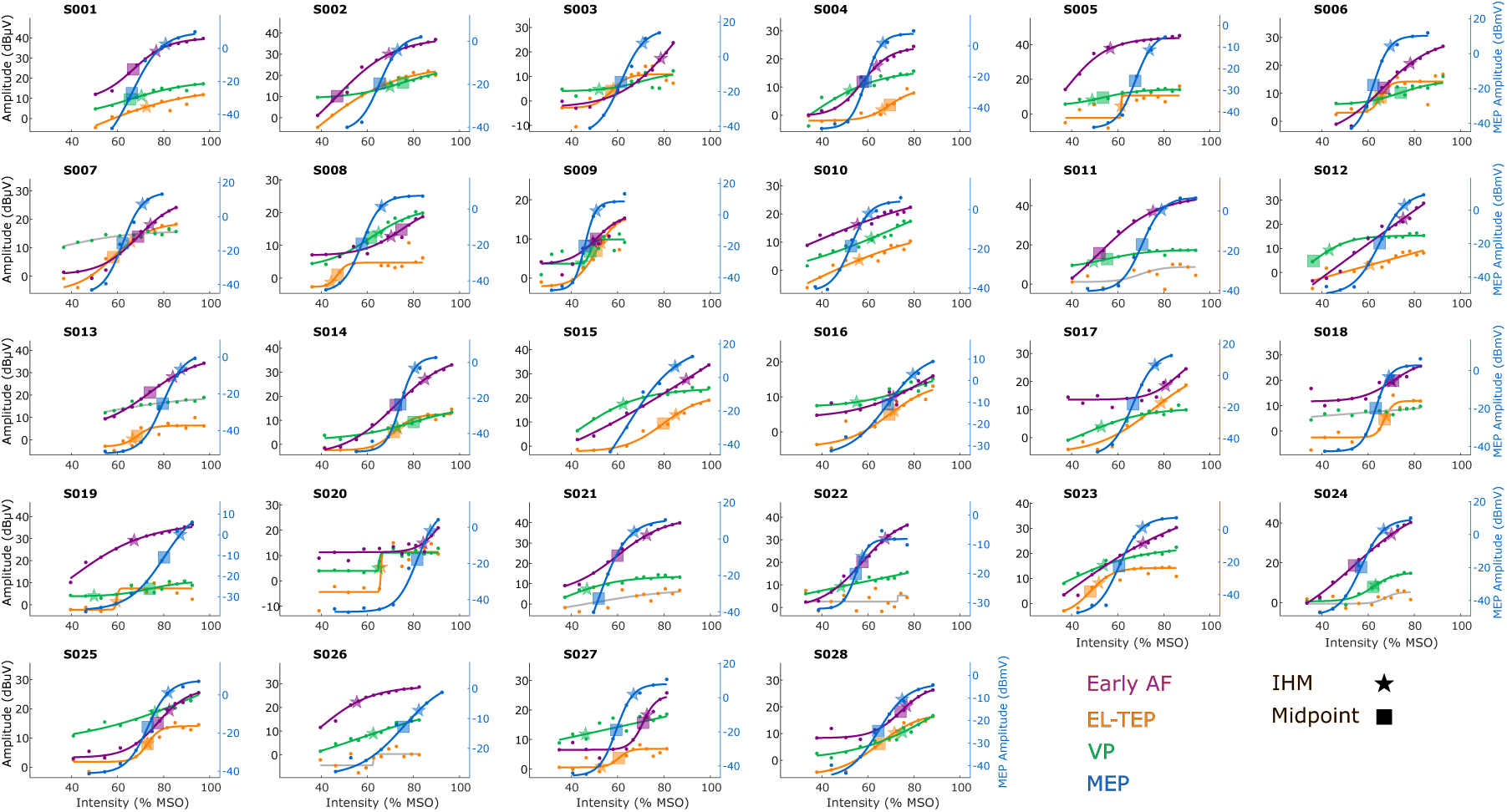
Individual I/O curves across response measures as a function of % MSO intensity. Individual input-output curves from 28 participants showing four TMS-evoked response measures plotted against stimulation intensity expressed as percentage of maximum stimulator output (% MSO). Purple curves and dots represent early artifact (AF, left y-axis), orange curves and dots represent EL-TEP amplitude (left y-axis), green curves and dots represent vertex potential (VP, left y-axis), and blue curves and dots represent motor evoked potential (MEP, right y-axis). Stars indicate intensity at half-maximum (IHM) values, and squares indicate sigmoid midpoints when calculable (see Section 2.6.6).

**Fig. S7.**
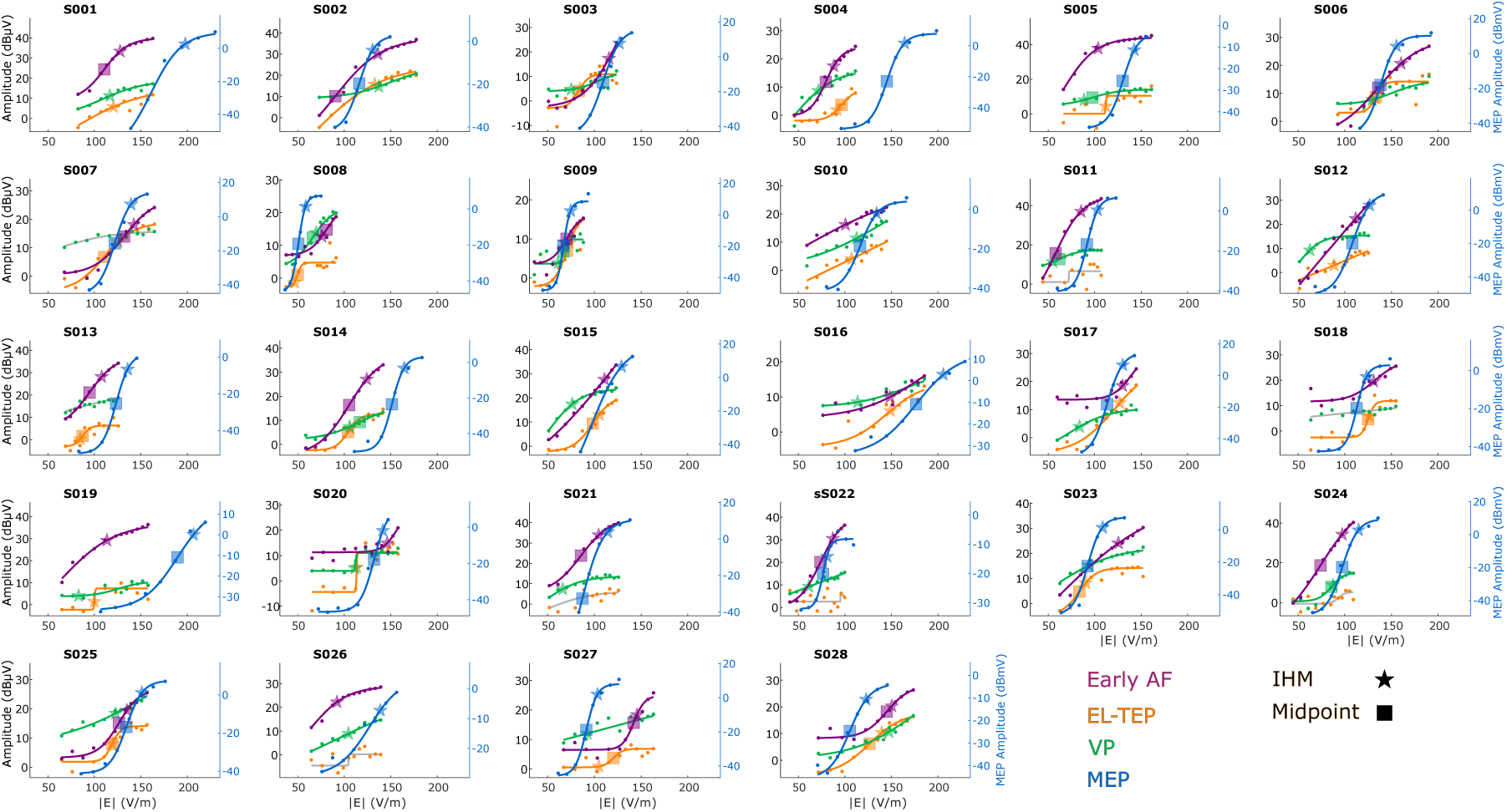
Individual I/O curves across response measures as a function of electric field magnitude (|E| V/m). Individual input-output curves from 28 participants showing four TMS-evoked response measures plotted against induced electric field magnitude at the cortical target (|E| in V/m). Purple curves and dots represent early artifact (AF, left y- axis), orange curves and dots represent EL-TEP amplitude (left y-axis), green curves and dots represent vertex potential (VP, left y-axis), and blue curves and dots represent motor evoked potential (MEP, right y-axis). Stars indicate intensity at half-maximum (IHM) values, and squares indicate sigmoid midpoints when calculable.

## References

[1] Miller EK, Cohen JD. An Integrative Theory of Prefrontal Cortex Function. Annu Rev Neurosci 2001;24:167–202. 10.1146/annurev.neuro.24.1.167.

[2] Etkin A, Büchel C, Gross JJ. The neural bases of emotion regulation. Nat Rev Neurosci 2015;16:693–700. 10.1038/nrn4044.

[3] Dixon ML, Thiruchselvam R, Todd R, Christoff K. Emotion and the prefrontal cortex: An integrative review. Psychol Bull 2017;143:1033–81. 10.1037/bul0000096.

[4] Friedman NP, Robbins TW. The role of prefrontal cortex in cognitive control and executive function. Neuropsychopharmacology 2022;47:72–89. 10.1038/s41386-021-01132-0.

[5] Polanía R, Nitsche MA, Ruff CC. Studying and modifying brain function with non-invasive brain stimulation. Nat Neurosci 2018;21:174–87. 10.1038/s41593-017-0054-4.

[6] Ressler KJ, Mayberg HS. Targeting abnormal neural circuits in mood and anxiety disorders: from the laboratory to the clinic. Nat Neurosci 2007;10:1116–24. 10.1038/nn1944.

[7] Drysdale AT, Grosenick L, Downar J, Dunlop K, Mansouri F, Meng Y, et al. Resting-state connectivity biomarkers define neurophysiological subtypes of depression. Nat Med 2016;23:nm.4246. 10.1038/nm.4246.

[8] Butt SJB, Lak A. The Spineless Origins of Prefrontal Cortex Dysfunction and Psychiatric Disorders. Neuron 2020;105:4–6. 10.1016/j.neuron.2019.12.009.

[9] Lefaucheur J-P, Aleman A, Baeken C, Benninger DH, Brunelin J, Di Lazzaro V, et al. Evidence-based guidelines on the therapeutic use of repetitive transcranial magnetic stimulation (rTMS): An update (2014–2018). Clin Neurophysiol 2020. 10.1016/j.clinph.2019.11.002.

[10] Cole EJ, Stimpson KH, Bentzley BS, Gulser M, Cherian K, Tischler C, et al. Stanford Accelerated Intelligent Neuromodulation Therapy for Treatment-Resistant Depression. Am J Psychiatry 2020;177:716–26. 10.1176/appi.ajp.2019.19070720.

[11] Vida RG, Sághy E, Bella R, Kovács S, Erdősi D, Józwiak-Hagymásy J, et al. Efficacy of repetitive transcranial magnetic stimulation (rTMS) adjunctive therapy for major depressive disorder (MDD) after two antidepressant treatment failures: meta-analysis of randomized sham-controlled trials. BMC Psychiatry 2023;23:545. 10.1186/s12888-023-05033-y.

[12] Gogulski J, Ross JM, Talbot A, Cline CC, Donati FL, Munot S, et al. Personalized Repetitive Transcranial Magnetic Stimulation for Depression. Biol Psychiatry Cogn Neurosci Neuroimaging 2023;8:351–60. 10.1016/j.bpsc.2022.10.006.

[13] Parmigiani S, Ross JM, Cline CC, Minasi CB, Gogulski J, Keller CJ. Reliability and Validity of Transcranial Magnetic Stimulation–Electroencephalography Biomarkers. Biol Psychiatry Cogn Neurosci Neuroimaging 2023;8:805–14. 10.1016/j.bpsc.2022.12.005.

[14] Kar SK. Predictors of Response to Repetitive Transcranial Magnetic Stimulation in Depression: A Review of Recent Updates. Clin Psychopharmacol Neurosci 2019;17:25–33. 10.9758/cpn.2019.17.1.25.

[15] Ross JM, Cline CC, Sarkar M, Truong J, Keller CJ. Neural effects of TMS trains on the human prefrontal cortex. Sci Rep 2023;13:22700. 10.1038/s41598-023-49250-7.

[16] Regenold WT, Deng Z-D, Lisanby SH. Noninvasive neuromodulation of the prefrontal cortex in mental health disorders. Neuropsychopharmacology 2022;47:361–72. 10.1038/s41386-021-01094-3.

[17] Koponen LM, Martinez M, Wood E, Murphy DLK, Goetz SM, Appelbaum LG, et al. Transcranial magnetic stimulation input–output curve slope differences suggest variation in recruitment across muscle representations in primary motor cortex. Front Hum Neurosci 2024;18.

[18] Krile L, Ensafi E, Cole J, Noor M, Protzner AB, McGirr A. A dose-response characterization of transcranial magnetic stimulation intensity and evoked potential amplitude in the dorsolateral prefrontal cortex. Sci Rep 2023;13:18650. 10.1038/s41598-023-45730-y.

[19] Carroll TJ, Riek S, Carson RG. Reliability of the input–output properties of the cortico- spinal pathway obtained from transcranial magnetic and electrical stimulation. J Neurosci Methods 2001;112:193–202. 10.1016/S0165-0270(01)00468-X.

[20] Therrien-Blanchet J-M, Ferland MC, Rousseau M-A, Badri M, Boucher E, Merabtine A, et al. Stability and test–retest reliability of neuronavigated TMS measures of corticospinal and intracortical excitability. Brain Res 2022;1794:148057. 10.1016/j.brainres.2022.148057.

[21] Devanne H, Lavoie BA, Capaday C. Input-output properties and gain changes in the human corticospinal pathway. Exp Brain Res 1997;114:329–38. 10.1007/PL00005641.

[22] Möller C, Arai N, Lücke J, Ziemann U. Hysteresis effects on the input–output curve of motor evoked potentials. Clin Neurophysiol 2009;120:1003–8. 10.1016/j.clinph.2009.03.001.

[23] Hill AT, Rogasch NC, Fitzgerald PB, Hoy KE. TMS-EEG: A window into the neurophysiological effects of transcranial electrical stimulation in non-motor brain regions. Neurosci Biobehav Rev 2016;64:175–84. 10.1016/j.neubiorev.2016.03.006.

[24] Farzan F, Vernet M, Shafi MMD, Rotenberg A, Daskalakis ZJ, Pascual-Leone A. Characterizing and Modulating Brain Circuitry through Transcranial Magnetic Stimulation Combined with Electroencephalography. Front Neural Circuits 2016;10. 10.3389/fncir.2016.00073.

[25] Farzan F. Transcranial Magnetic Stimulation–Electroencephalography for Biomarker Discovery in Psychiatry. Biol Psychiatry 2024;95:564–80. 10.1016/j.biopsych.2023.12.018.

[26] Gogulski J, Cline CC, Ross JM, Truong J, Sarkar M, Parmigiani S, et al. Mapping cortical excitability in the human dorsolateral prefrontal cortex. Clin Neurophysiol 2024;164:138–48. 10.1016/j.clinph.2024.05.008.

[27] Gogulski J, Cline CC, Ross JM, Parmigiani S, Keller CJ. Reliability of the TMS-evoked potential in dorsolateral prefrontal cortex. Cereb Cortex 2024;34:bhae130. 10.1093/cercor/bhae130.

[28] Massimini M, Ferrarelli F, Huber R, Esser SK, Singh H, Tononi G. Breakdown of Cortical Effective Connectivity During Sleep. Science 2005;309:2228–32. 10.1126/science.1117256.

[29] Massimini M, Ferrarelli F, Esser SK, Riedner BA, Huber R, Murphy M, et al. Triggering sleep slow waves by transcranial magnetic stimulation. Proc Natl Acad Sci 2007;104:8496–501. 10.1073/pnas.0702495104.

[30] Rosanova M, Casali A, Bellina V, Resta F, Mariotti M, Massimini M. Natural Frequencies of Human Corticothalamic Circuits. J Neurosci 2009;29:7679–85. 10.1523/JNEUROSCI.0445-09.2009.

[31] Ross JM, Sarkar M, Keller CJ. Experimental suppression of transcranial magnetic stimulation-electroencephalography sensory potentials. Hum Brain Mapp 2022;n/a. 10.1002/hbm.25990.

[32] Casarotto S, Fecchio M, Rosanova M, Varone G, D’Ambrosio S, Sarasso S, et al. The rt-TEP tool: real-time visualization of TMS-Evoked Potentials to maximize cortical activation and minimize artifacts. J Neurosci Methods 2022;370:109486. 10.1016/j.jneumeth.2022.109486.

[33] Parmigiani S, Cline CC, Sarkar M, Forman L, Truong J, Ross JM, et al. Real-time optimization to enhance noninvasive cortical excitability assessment in the human dorsolateral prefrontal cortex. Clin Neurophysiol 2025:S1388245725003049. 10.1016/j.clinph.2025.02.261.

[34] Cline CC, Forman L, Hartford W, Truong J, Parmigiani S, Keller CJ. NaviNIBS: a comprehensive and open-source software toolbox for neuronavigated noninvasive brain stimulation. J Neural Eng 2025;22:056007. 10.1088/1741-2552/adfab2.

[35] Lasso A, Heffter T, Rankin A, Pinter C, Ungi T, Fichtinger G. PLUS: Open-Source Toolkit for Ultrasound-Guided Intervention Systems. IEEE Trans Biomed Eng 2014;61:2527–37. 10.1109/TBME.2014.2322864.

[36] Tokuda J, Fischer GS, Papademetris X, Yaniv Z, Ibanez L, Cheng P, et al. OpenIGTLink: an open network protocol for image-guided therapy environment. Int J Med Robot 2009;5:423–34. 10.1002/rcs.274.

[37] Thielscher A, Antunes A, Saturnino GB. Field modeling for transcranial magnetic stimulation: A useful tool to understand the physiological effects of TMS? 2015 37th Annu. Int. Conf. IEEE Eng. Med. Biol. Soc. EMBC, 2015, p. 222–5. 10.1109/EMBC.2015.7318340.

[38] Nielsen JD, Madsen KH, Puonti O, Siebner HR, Bauer C, Madsen CG, et al. Automatic skull segmentation from MR images for realistic volume conductor models of the head: Assessment of the state-of-the-art. NeuroImage 2018;174:587–98. 10.1016/j.neuroimage.2018.03.001.

[39] Julkunen P, Säisänen L, Hukkanen T, Danner N, Könönen M. Does second-scale intertrial interval affect motor evoked potentials induced by single-pulse transcranial magnetic stimulation? Brain Stimulat 2012;5:526–32. 10.1016/j.brs.2011.07.006.

[40] Cline CC, Lucas MV, Sun Y, Menezes M, Etkin A. Advanced Artifact Removal for Automated TMS-EEG Data Processing. 2021 10th Int. IEEEEMBS Conf. Neural Eng. NER, 2021, p. 1039–42. 10.1109/NER49283.2021.9441147.

[41] Pion-Tonachini L, Kreutz-Delgado K, Makeig S. ICLabel: An automated electroencephalographic independent component classifier, dataset, and website. NeuroImage 2019;198:181–97. 10.1016/j.neuroimage.2019.05.026.

[42] Mutanen TP, Metsomaa J, Liljander S, Ilmoniemi RJ. Automatic and robust noise suppression in EEG and MEG: The SOUND algorithm. NeuroImage 2018;166:135–51. 10.1016/j.neuroimage.2017.10.021.

[43] Mutanen TP, Biabani M, Sarvas J, Ilmoniemi RJ, Rogasch NC. Source-based artifact-rejection techniques available in TESA, an open-source TMS–EEG toolbox. Brain Stimulat 2020;13:1349–51. 10.1016/j.brs.2020.06.079.

[44] Huang Y, Hajnal B, Entz L, Fabó D, Herrero JL, Mehta AD, et al. Intracortical Dynamics Underlying Repetitive Stimulation Predicts Changes in Network Connectivity. J Neurosci 2019;39:6122–35. 10.1523/JNEUROSCI.0535-19.2019.

[45] Keller CJ, Huang Y, Herrero JL, Fini ME, Du V, Lado FA, et al. Induction and Quantification of Excitability Changes in Human Cortical Networks. J Neurosci 2018;38:5384–98. 10.1523/JNEUROSCI.1088-17.2018.

[46] Bergmann TO, Mölle M, Schmidt MA, Lindner C, Marshall L, Born J, et al. EEG-Guided Transcranial Magnetic Stimulation Reveals Rapid Shifts in Motor Cortical Excitability during the Human Sleep Slow Oscillation. J Neurosci 2012;32:243–53. 10.1523/JNEUROSCI.4792-11.2012.

[47] Casarotto S, Canali P, Rosanova M, Pigorini A, Fecchio M, Mariotti M, et al. Assessing the Effects of Electroconvulsive Therapy on Cortical Excitability by Means of Transcranial Magnetic Stimulation and Electroencephalography. Brain Topogr 2013;26:326–37. 10.1007/s10548-012-0256-8.

[48] Gosseries O, Thibaut A, Boly M, Rosanova M, Massimini M, Laureys S. Assessing consciousness in coma and related states using transcranial magnetic stimulation combined with electroencephalography. Ann Fr Anesth Réanimation 2014;33:65–71. 10.1016/j.annfar.2013.11.002.

[49] Eshel N, Keller CJ, Wu W, Jiang J, Mills-Finnerty C, Huemer J, et al. Global connectivity and local excitability changes underlie antidepressant effects of repetitive transcranial magnetic stimulation. Neuropsychopharmacology 2020;45:1018–25. 10.1038/s41386-020-0633-z.

[50] Voineskos D, Blumberger DM, Zomorrodi R, Rogasch NC, Farzan F, Foussias G, et al. Altered Transcranial Magnetic Stimulation-Electroencephalographic Markers of Inhibition and Excitation in the Dorsolateral Prefrontal Cortex in Major Depressive Disorder. Biol Psychiatry 2018. 10.1016/j.biopsych.2018.09.032.

[51] Hassan U, Pillen S, Zrenner C, Bergmann TO. The Brain Electrophysiological recording & STimulation (BEST) toolbox. Brain Stimulat 2022;15:109–15. 10.1016/j.brs.2021.11.017.

[52] Michel CM, Murray MM. Towards the utilization of EEG as a brain imaging tool. NeuroImage 2012;61:371–85. 10.1016/j.neuroimage.2011.12.039.

[53] Gramfort A, Papadopoulo T, Olivi E, Clerc M. OpenMEEG: opensource software for quasistatic bioelectromagnetics. Biomed Eng OnLine 2010;9:45. 10.1186/1475-925X-9-45.

[54] Kybic J, Clerc M, Abboud T, Faugeras O, Keriven R, Papadopoulo T. A common formalism for the Integral formulations of the forward EEG problem. IEEE Trans Med Imaging 2005;24:12–28. 10.1109/TMI.2004.837363.

[55] Tadel F, Baillet S, Mosher JC, Pantazis D, Leahy RM. Brainstorm: A User-Friendly Application for MEG/EEG Analysis. Comput Intell Neurosci 2011;2011. 10.1155/2011/879716.

[56] Bates D, Mächler M, Bolker B, Walker S. Fitting Linear Mixed-Effects Models Using lme4. J Stat Softw 2015;67:1–48. 10.18637/jss.v067.i01.

[57] Luke SG. Evaluating significance in linear mixed-effects models in R. Behav Res Methods 2017;49:1494–502. 10.3758/s13428-016-0809-y.

[58] Lenth RV, Banfai B, Bolker B, Buerkner P, Giné-Vázquez I, Hervé M, et al. emmeans: Estimated Marginal Means, aka Least-Squares Means 2025.

[59] Lin LI-K. A Concordance Correlation Coefficient to Evaluate Reproducibility. Biometrics 1989;45:255–68. 10.2307/2532051.

[60] Kerwin LJ, Keller CJ, Wu W, Narayan M, Etkin A. Test-retest reliability of transcranial magnetic stimulation EEG evoked potentials. Brain Stimul Basic Transl Clin Res Neuromodulation 2017;0. 10.1016/j.brs.2017.12.010.

[61] Caulfield KA, Li X, George MS. Four electric field modeling methods of Dosing Prefrontal Transcranial Magnetic Stimulation (TMS): Introducing APEX MT dosimetry. Brain Stimul Basic Transl Clin Res Neuromodulation 2021;14:1032–4. 10.1016/j.brs.2021.06.012.

[62] Desai NS, Rutherford LC, Turrigiano GG. Plasticity in the intrinsic excitability of cortical pyramidal neurons. Nat Neurosci 1999;2:515–20. 10.1038/9165.

[63] Marom S, Marder E. A biophysical perspective on the resilience of neuronal excitability across timescales. Nat Rev Neurosci 2023;24:640–52. 10.1038/s41583-023-00730-9.

[64] Fox KCR, Shi L, Baek S, Raccah O, Foster BL, Saha S, et al. Intrinsic network architecture predicts the effects elicited by intracranial electrical stimulation of the human brain. Nat Hum Behav 2020;4:1039–52. 10.1038/s41562-020-0910-1.

[65] Yun R, Mishler JH, Perlmutter SI, Rao RPN, Fetz EE. Responses of Cortical Neurons to Intracortical Microstimulation in Awake Primates. eNeuro 2023;10. 10.1523/ENEURO.0336-22.2023.

[66] Sukenik N, Vinogradov O, Weinreb E, Segal M, Levina A, Moses E. Neuronal circuits overcome imbalance in excitation and inhibition by adjusting connection numbers. Proc Natl Acad Sci 2021;118:e2018459118. 10.1073/pnas.2018459118.

[67] Remme MWH, Wadman WJ. Homeostatic Scaling of Excitability in Recurrent Neural Networks. PLOS Comput Biol 2012;8:e1002494. 10.1371/journal.pcbi.1002494.

[68] Lioumis P, Kičić D, Savolainen P, Mäkelä JP, Kähkönen S. Reproducibility of TMS— Evoked EEG responses. Hum Brain Mapp 2009;30:1387–96. 10.1002/hbm.20608.

[69] Elston GN. Cortex, Cognition and the Cell: New Insights into the Pyramidal Neuron and Prefrontal Function. Cereb Cortex 2003;13:1124–38. 10.1093/cercor/bhg093.

[70] Douglas RJ, Martin KAC. Neuronal circuits of the neocortex. Annu Rev Neurosci 2004;27:419–51. 10.1146/annurev.neuro.27.070203.144152.

[71] Cash RFH, Noda Y, Zomorrodi R, Radhu N, Farzan F, Rajji TK, et al. Characterization of Glutamatergic and GABAA-Mediated Neurotransmission in Motor and Dorsolateral Prefrontal Cortex Using Paired-Pulse TMS–EEG. Neuropsychopharmacology 2017;42:502–11. 10.1038/npp.2016.133.

[72] Godlove DC, Maier A, Woodman GF, Schall JD. Microcircuitry of agranular frontal cortex: testing the generality of the canonical cortical microcircuit. J Neurosci 2014;34:5355–69.

[73] Weise K, Makaroff SN, Numssen O, Bikson M, Knösche TR. Statistical method accounts for microscopic electric field distortions around neurons when simulating activation thresholds. Brain Stimulat 2025. 10.1016/j.brs.2025.02.007.

[74] Numssen O, Kuhnke P, Weise K, Hartwigsen G. Electric field based dosing for TMS. Imaging Neurosci 2024. 10.1162/imag_a_00106.

[75] Dannhauer M, Gomez LJ, Robins PL, Wang D, Hasan NI, Thielscher A, et al. Electric Field Modeling in Personalizing Transcranial Magnetic Stimulation Interventions. Biol Psychiatry 2024;95:494–501. 10.1016/j.biopsych.2023.11.022.

[76] Czerwonky DM, Aberra AS, Gomez LJ. A boundary element method of bidomain modeling for predicting cellular responses to electromagnetic fields. J Neural Eng 2024;21:036050. 10.1088/1741-2552/ad5704.

[77] Siebner HR, Funke K, Aberra AS, Antal A, Bestmann S, Chen R, et al. Transcranial magnetic stimulation of the brain: What is stimulated? – A consensus and critical position paper. Clin Neurophysiol 2022;140:59–97. 10.1016/j.clinph.2022.04.022.

[78] Wang JB, Hassan U, Bruss JE, Oya H, Uitermarkt BD, Trapp NT, et al. Effects of transcranial magnetic stimulation on the human brain recorded with intracranial electrocorticography. Mol Psychiatry 2024;29:1228–40. 10.1038/s41380-024-02405-y.

[79] Michel CM, Murray MM, Lantz G, Gonzalez S, Spinelli L, Grave de Peralta R. EEG source imaging. Clin Neurophysiol 2004;115:2195–222. 10.1016/j.clinph.2004.06.001.

[80] He B, Sohrabpour A, Brown E, Liu Z. Electrophysiological Source Imaging: A Noninvasive Window to Brain Dynamics. Annu Rev Biomed Eng 2018;20:171–96. 10.1146/annurev-bioeng-062117-120853.

[81] Subramanian AK, Talbot A, Kim N, Parmigiani S, Cline CC, Solomon EA, et al. Scalp EEG predicts intracranial brain activity in humans 2025:2025.04.07.647612. 10.1101/2025.04.07.647612.

[82] Pigorini A, Avanzini P, Barborica A, Bénar C-G, David O, Farisco M, et al. Simultaneous invasive and non-invasive recordings in humans: A novel Rosetta stone for deciphering brain activity. J Neurosci Methods 2024;408:110160. 10.1016/j.jneumeth.2024.110160.

[83] Briley PM, Webster L, Lankappa S, Pszczolkowski S, McAllister-Williams RH, Liddle PF, et al. Trajectories of improvement with repetitive transcranial magnetic stimulation for treatment-resistant major depression in the BRIGhTMIND trial. Npj Ment Health Res 2024;3:32. 10.1038/s44184-024-00077-8.

[84] Lacroix A, Calvet B, Laplace B, Lannaud M, Plansont B, Guignandon S, et al. Predictors of clinical response after rTMS treatment of patients suffering from drug- resistant depression. Transl Psychiatry 2021;11:587. 10.1038/s41398-021-01555-9.

[85] Fox MD, Buckner RL, White MP, Greicius MD, Pascual-Leone A. Efficacy of Transcranial Magnetic Stimulation Targets for Depression Is Related to Intrinsic Functional Connectivity with the Subgenual Cingulate. Biol Psychiatry 2012;72:595– 603. 10.1016/j.biopsych.2012.04.028.

[86] Jung J, Ralph MAL, Jackson RL. Subregions of DLPFC Display Graded yet Distinct Structural and Functional Connectivity. J Neurosci 2022;42:3241–52. 10.1523/JNEUROSCI.1216-21.2022.

[87] Fox MD, Liu H, Pascual-Leone A. Identification of reproducible individualized targets for treatment of depression with TMS based on intrinsic connectivity. NeuroImage 2013;66:151–60. 10.1016/j.neuroimage.2012.10.082.

[88] Solomon EA, Wang JB, Oya H, Howard MA, Trapp NT, Uitermarkt BD, et al. TMS provokes target-dependent intracranial rhythms across human cortical and subcortical sites. Brain Stimulat 2024;17:698–712. 10.1016/j.brs.2024.05.014.

[89] Li Z, Liu X, Tatz J, Hassan U, Wang JB, Keller CJ, et al. A Practical Preprocessing Pipeline for Concurrent TMS-iEEG: Critical Steps and Methodological Considerations 2025:2025.08.13.670238. 10.1101/2025.08.13.670238.

[90] Huang Y, Zelmann R, Hadar P, Dezha-Peralta J, Richardson RM, Williams ZM, et al. Theta-burst direct electrical stimulation remodels human brain networks. Nat Commun 2024;15:6982. 10.1038/s41467-024-51443-1.

